# Structural Mechanism of Prestin-Membrane Mechanotransduction

**DOI:** 10.1101/2025.10.31.685930

**Authors:** Navid Bavi, Patrick R Haller, Kazuaki Homma, Xiaoxuan Lin, Wieslawa Milewski, Minglei Zhao, Tobin Sosnick, Eduardo Perozo

## Abstract

Sound frequency discrimination in mammals depends on the conformational transitions of prestin (SLC26A5), the piezoelectric motor in outer hair cells. The mechanism that enables prestin’s electrically driven interconversion and its dependence on membrane mechanics, remains unresolved. Here, we show that membrane forces represent a strong driver of the same conformational changes generated by transmembrane voltage and stabilized by bound anions. Single particle cryo-EM structures of nanodisc-reconstituted prestin were obtained from varying lipid composition and membrane thickness. These structures show that membrane thinning biases prestin from a compact conformation to a fully expanded conformation, mimicking outer hair cell elongation/contraction during electromotility. In contrast, zebrafish SLC26A5 transporters undergo complete elevator movements with redistribution of areal changes across leaflets. The structures, together with mutagenesis, H/D exchange mass spectrometry data, and NLC measurements, offer a high-resolution understanding of how prestin translates membrane tension into charge and motor movement during sound-evoked vibrations, revealing a process of reciprocal electro-mechanical transduction essential for tuning cochlear amplification.

## Introduction

Hearing in mammals relies on the electromechanical activity of outer hair cells (OHCs), which elongate and shorten in response to changes in membrane voltage to amplify sound within the cochlea (*1, 2*). This voltage-driven mechanical response, known as electromotility, is powered by prestin (SLC26A5) (*3*). Unlike every other members of the SLC26 family (*4–9*), prestin has evolved not as a transporter but as a piezoelectric membrane motor (*10–14*), originally thought to populate two fundamental conformations with different cross sectional areas in the plane of the bilayer (*15*). Despite recent advances on the structural determination of SLC26 proteins (*16–28*), the molecular basis for this unique specialization and its dependence on lipid bilayer mechanics remains enigmatic.

Prestin’s nonlinear capacitance (NLC) serves as a hallmark of its electromechanical activity, reflecting voltage-dependent transitions between its conformational states (*29, 30*). The binding of anions, such as Cl^-^and salicylate, significantly influences NLC, underscoring the importance of anion interactions in prestin’s function (*31, 32*). While recent high-resolution structures of have provided mechanistic insights into the influence of anion binding and their influence on in-plane areal expansion (*16, 18*), the precise molecular basis by which mechanical forces influence its charge-transfer properties (*33*) and the dynamics of bound anions remains obscure.

In this study, we demonstrate how prestin directly responds to mechanical forces according to the “force-from-lipids” principle (*34–36*), as it has been observed in mechanosensitive ion channels. Using single-particle cryo-electron microscopy (cryo-EM) in physiological Cl^-^, we resolved eight distinct structures to systematically dissect prestin’s conformational cycle under different membrane forces. Together with functional assays, and hydrogen-deuterium exchange mass spectrometry (HDX-MS), we identified prestin’s gate domain (TMs 6-7) as a key mechanosensory element and elucidate how the lipid environment modulates occupancy of prestin’s mechanical and electrical states. Furthermore, by solving the structures of non-motor zebrafish prestin in multiple conformations and in nanodiscs of varying thickness, we contrast how SLC26A5 transporters respond to membrane forces compared with a motor behavior. These data illuminate the functional and structural adaptations that distinguish electromotile prestin from its transporter relatives, providing mechanistic insight into the evolution of cochlear amplification.

## Results

### Membrane tension and bilayer thickness tunes the operating voltage of prestin

It has been shown that prestin’s NLC response is sensitive to membrane tension (*37–39*) and membrane thickness (*11, 40*) within the area expansion model (*15*). First, we confirmed, dolphin prestin (*41, 42*) sensitivity to changes in membrane properties by pressure-clamp electrophysiology on overexpressing HEK293 cells, with control recordings were carried out on native outer hair cells (OHCs) isolated from adult mice (**fig. 1**). Increases in intracellular pressure, expected to augment membrane tension, caused depolarizing shifts in the voltage at peak capacitance (V_pk_) of NLC in both OHCs and HEK293 cells (**fig. 1, A and B)**. Specifically, V_pk_ shifted at a rate of +6.9 ± 0.3 mV/mmHg when applying increasing pressure on OHCs. Similarly, 10 mmHg pressure on HEK293 cells overexpressing dolphin prestin shifted the V_pk_ from −57.3 ± 4.8 mV to −31.1 ± 11.1 mV. Pressure-induced changes also increased linear capacitance (C_lin_) in most HEK293 cells, due to nontrivial cell swelling under positive turgor. These findings are in agreement with previous reports (*37–39, 43*), and confirm that dolphin prestin functions as an intrinsic mechanosensor, detecting and adapting to membrane tension in both native and heterologous systems.

**Fig. 1:**
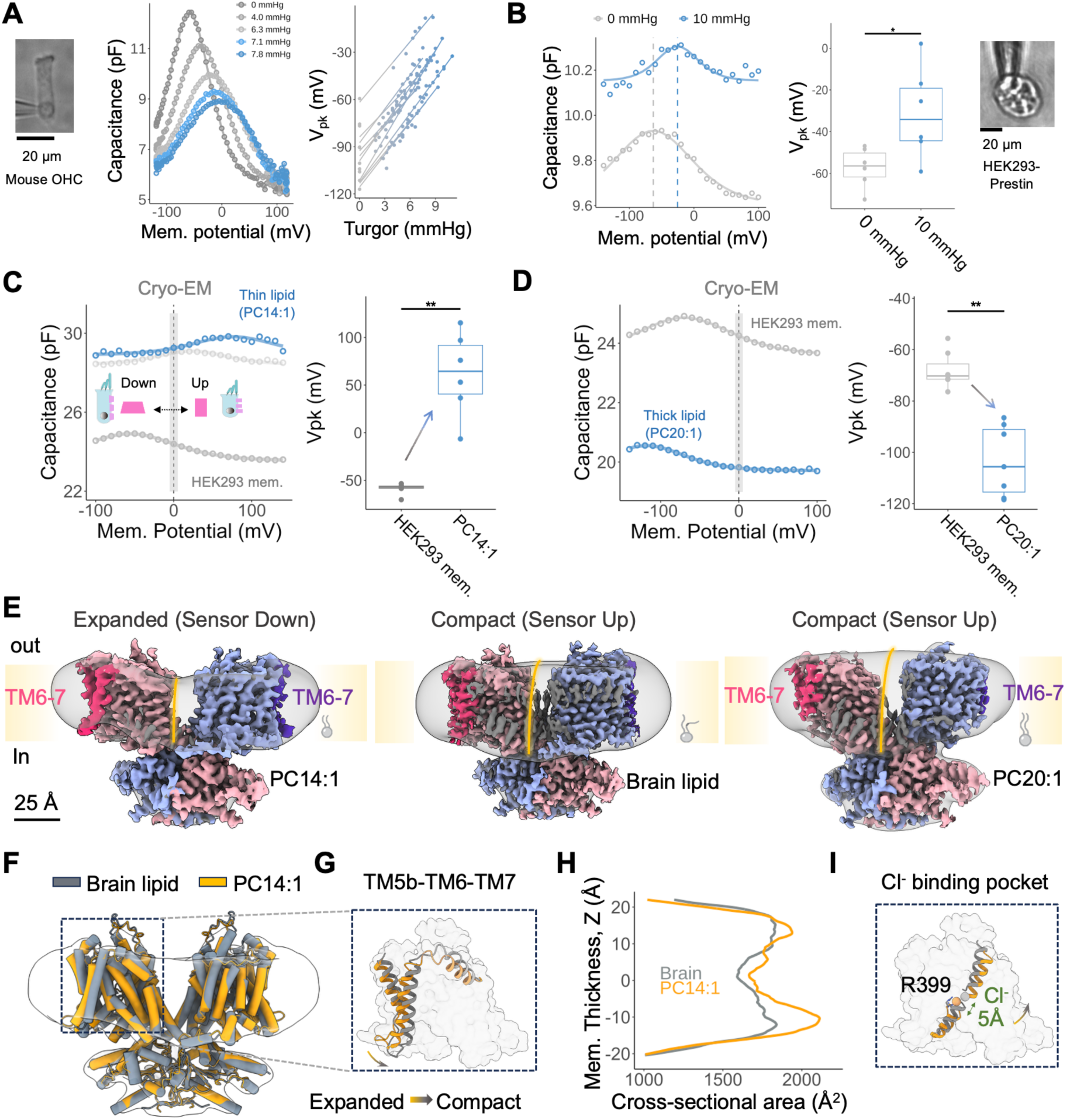
Prestin’s conformational cycle is tightly linked to membrane mechanics. A) Whole-cell patch-clamp electrophysiology was performed on outer hair cells isolated from adult (P21) mice and on B) HEK293 cells overexpressing dolphin prestin. Both were subjected to controlled intracellular pressure to induce plasma membrane tension. This approach mimicked outer hair cell elongation and shrinkage, enabling the monitoring of its effect on prestin’s charge transfer. In both cell types, the NLC exhibited depolarizing shifts with increasing tension: A) For outer hair cells, the V_pk_ shifted at a rate of +6.9 ± 0.3 mV/mmHg (n = 10; mean ± SEM). B) For HEK293 cells, V_pk_ shifted from −57.3 ± 4.8 mV to −31.1 ± 11.1 mV (n = 6; mean ± SEM) after application of 10 mmHg pressure. Linear capacitance slightly increased in most cases for HEK293 cells, likely due to membrane swelling. These results demonstrate the mechanosensitivity of prestin’s charge transfer in both native and heterologous systems. C) Whole-cell patch-clamp electrophysiology was performed on HEK293 cells transfected with wild-type dolphin prestin. Nonlinear capacitance (NLC) was recorded before and after the addition of 1 mM α-cyclodextrin doped with PC14:1 (thin) lipids, mimicking membrane thinning during somatic elongation. Lipid delivery shifted the V_pk_ of NLC from −57.8 ± 3.0 mV to depolarized potentials (+61.9 ± 21.0 mV; mean ± SEM), accompanied by an increase in linear capacitance, likely due to reduced membrane thickness. D) To simulate outer hair cell contraction, α-cyclodextrin was doped with PC20:1 (thick) lipids and delivered to the plasma membrane of HEK293 cells transfected with dolphin prestin. This caused a hyperpolarizing shift of V_pk_ from −57.8 ± 3.0 mV to −93.6 ± 4.2 mV and a decrease in linear capacitance, consistent with membrane thickening. Together, these results demonstrate that membrane thickness modulates charge transfer in prestin, transitioning between the contracted (sensor up) and expanded (sensor down) states (n = 6; mean ± SEM). E) Cryo-EM density maps and the overall structure of the dolphin prestin homodimer solved in MSP-E3D1 nanodiscs under three lipid conditions: PC14:1 (left, expanded state), brain lipid extract (middle, compact state), and PC20:1 (right, compact state). The details of nanodiscs composition is explained in the Methods. The nominal resolutions of the structures are 3.8 Å, 2.9 Å, and 3.1 Å, respectively. Scale bar, 25 Å. Yellow lines indicate nanodisc thickness. Monomers are shown in blue and salmon. The TM6-7 region is highlighted in dark pink and dark violet. Note that the TM6-7 region in the PC20:1 structure is not fully resolved, while the TM6-7 helices in the PC14:1 lipid (thin lipid) exhibit outward tilting. F) Structural overlay of prestin in the compact and expanded states, depicted as cartoon models. G) Transmembrane helices TM5b-TM6-TM7 undergo significant conformational changes, bending at the gate region to accommodate varying bilayer thickness. H) Membrane thinning induces intra-membrane areal-expansion of prestin. Residues 460 to 505 were used for dimer alignment, and cross-sectional area changes were calculated for a single monomer. I) Anion density, previously identified as Cl⁻, was observed within the binding site in both the contracted and expanded states, suggesting its role in the electromotive cycle. As the binding-site undergoes an elevator-like motion, Cl⁻ elevates ∼5 Å from the expanded to the compact conformation.

To correlate membrane thinning effects with tension, we modulated bilayer thickness using α-cyclodextrin, to facilitates lipid exchange with cellular membranes (see Methods). Incorporation of phospholipids with short acyl chains (PC14:1; thin lipids) induced a depolarizing shift in the V_pk_ from - 58.7 ± 3.0 mV to +61.9 ± 21.0 mV, mimicking membrane thinning during somatic elongation of OHCs (**fig. 1, C and D**). Conversely, incorporation of phospholipids with long acyl chains (PC20:1; thick lipids), caused a hyperpolarizing shift in V_pk_ to −93.6 ± 4.2, corresponding to membrane thickening during OHC contraction. These shifts in V_pk_ were accompanied by changes in linear capacitance, as expected if membrane thickness had a direct influence on prestin’s charge transfer and electromotility (*11, 40*).

### Membrane thinning switches Prestin from compact to the expanded conformation in Cl⁻

We used membrane thinning and “lipid hijacking” by undoped α-cyclodextrin (*11, 44, 45*) as a structural proxy for the membrane forces exerted during OHCs’ electromotility. Although conceptually distinct, these approaches effectively mimic the biophysical effects of tension and enable high-resolution cryo-EM studies of prestin under mechanically relevant conditions. Prestin, was reconstituted into nanodiscs of different lipid compositions and in the presence of Cl^-^ using an “on-column” reconstitution method (*46, 47*). Nanodiscs structures (**fig. 1, figs. S1 to S5)** were obtained under three lipid conditions: thin lipids (PC14:1, at 3.5 Å), “mixed/regular lipids” (porcine brain lipid; at 2.9 Å), and thick lipids (PC20:1, at 3.0 Å) (**fig. 1E; figs. S2 to S5**). Similar to the previously reported structures in detergent (*16–18, 28*), prestin is observed as a symmetric homodimer (C2 symmetry) (**fig. 1F, figs. S2 to S4**). The 14 TM region is organized into two distinct domains that are a characteristic of SLC26 and other elevator-type transporters (*20–22, 27, 48*): a gate domain, consisting of TM segments 5–7 and 12–14, and a core domain, which includes TM segments 1–4 and 8–11. Transmembrane segments are domain-swapped with the cytoplasmic region, and both the N- and C-termini, including the sulfate transporter and anti-sigma factor antagonist (STAS) domain, are positioned within the cytoplasmic space (**fig. S4**).

Under thin lipid conditions (PC14:1), prestin predominantly adopted the expanded conformation, characterized by the movement of the anion binding site towards the cytoplasm (“Sensor Down”), and a pronounced bending and tilting of the peripheral transmembrane helices 6 and 7 (TM6-7), known as the “electromotility elbow” (*16*) (**fig. 1, F-I**). The bending motion of TM6-7 was further accompanied by more subtle movements in the gate domain, particularly in TM5b, to accommodate hydrophobic mismatch with the bilayer (**fig. 1G).** In contrast, thick lipids (PC20:1) and mixed brain lipid environments stabilized a compact “Sensor Up” state. Notably, the brain lipid structure resembled the Cl⁻-bound state observed in thick lipid nanodiscs, although the TM6-7 region exhibited putative high flexibility/dynamics as seen by unresolved density in thick lipid conditions (**fig. S5D**). These structural transitions suggest a crucial role of the gate domain as a mechanosensitive region (in particular TM5-7), bending and expanding to adapt to membrane deformation. Area changes (per monomer) of up to ∼150 Å² (∼10 %) across the membrane were observed in the inner leaflet, with alterations in membrane thickness (**fig. 1H**). These area changes are similar to the ones observed in anion-induced conformational transitions (*16, 18*) **(fig. S5E)**. We note that anion-binding interactions were preserved in all conditions, and well-resolved Cl⁻ densities were observed in both the expanded and compact states. In accordance with elevator-like movements at the core-domain, the Cl⁻ density shifted upward by approximately 5 Å when transitioning from the expanded to the compact state (**fig. 1I**). However, no difference was observed between “Sensor Down” vs. “Sensor Up” conformations for the region separating the TM3-TM10 helical dipoles (distance *d*) (**fig. S5, F and G**). This contrasts with the differences in binding site volumes in the presence of different anions seen in previous studies (*16, 18*).

### Single glycine substitutions push Prestin across its entire electromotility cycle

Using patch-clamp electrophysiology, we identified a critical role for glycine residues on TM6 in modulating the NLC in prestin (*16*). Specifically, mutation G274I in TM6 induces subtle hyperpolarizing shifts, while G275I abolishes NLC in the measurable voltage range. The structural basis of these functional changes was analyzed from the cryo-EM structures of G274I and G275I in detergent (at 2.9 Å and 3.4 Å, respectively) (**fig. 2C, fig. S7**). Consistent with the NLC measurements, G274I adopted a compact conformation with its elevator in the Up state, whereas the G275I mutant exhibited an expanded conformation similar to the elevator Down state (**fig. 2, D and E**). Structural overlays revealed that introduction of an isoleucine at these positions altered the bending motion of the TM6-7 helix (**fig. 2C).** So, while TM6 appears straightened at the hinge in the G274I mutant, the G275I mutation induced a kink located above its native position in wild-type prestin. Furthermore, membrane area changes associated with G274I and G275I aligned with lipid-induced thinning experiments. This highlights the tight coupling between prestin’s structural states and bilayer mechanics (**fig. 2D**). Notably, despite these structural perturbations, the anion-binding site remained Cl- occupied in both G274I and G275I mutants, consistent with WT prestin structures in lipid nanodiscs (**fig. 2E**). This finding suggests that while glycine mutations disrupt the flexibility of TM6-7, altering prestin’s conformational states, they don’t seem to affect the anion-binding site, as expected from the role of this motif in prestin’s electromotive cycle.

**Fig. 2:**
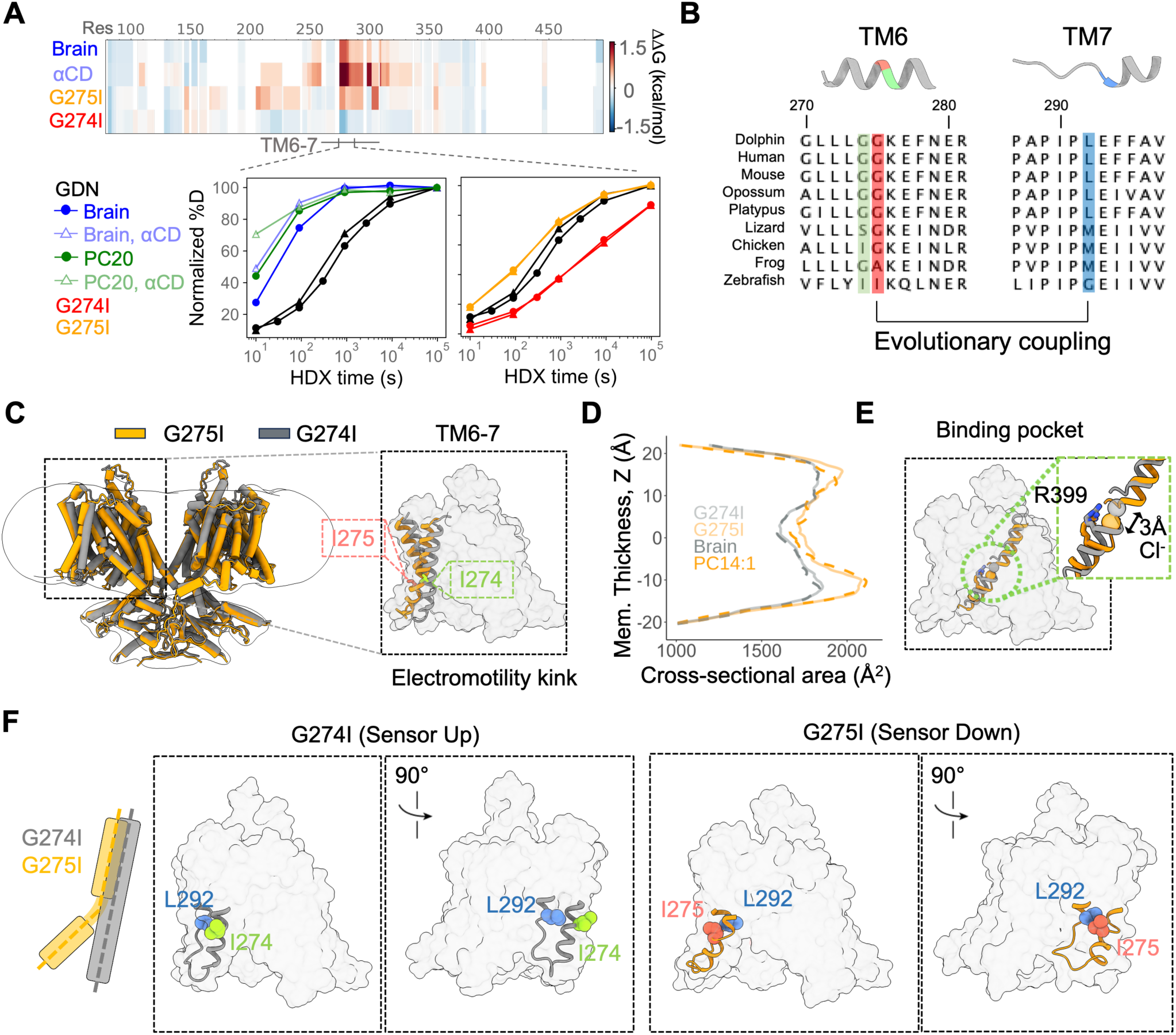
The role of the evolutionarily conserved flexible region in the TM6-7 gate domain in sensing membrane-mediated mechanical forces. A) The TM6-7 region in the gate domain, which is lipid-facing, showed the most pronounced conformational changes in response to alterations in the lipid environment. This was confirmed using structural studies under various conditions, including mutagenesis, lipid delivery via α-cyclodextrin doped thick (PC20:1) lipids, and lipid extraction using empty α-cyclodextrin. Hydrogen-deuterium exchange mass spectrometry (HDX-MS) revealed that the TM6-7 region, particularly after the double Gly274-Gly275 residues, exhibited the greatest structural adaptability to changes in membrane properties. These residues are evolutionarily conserved, and mutations in this region severely impact charge transfer. B) Sequence alignment of TM6 (residues 270-281) and TM7 (residues 287-297) helices, highlighting that glycine residues 274 and 275 on TM6 are unique to mammals, and illustrating the coupling between residues 275 on TM6 and 292 on TM7. C) Structure overlay of prestin mutants G274I and G275I depicted as cartoon. Structures were determined in the presence of Cl⁻ and in detergent at nominal resolutions of 2.9A and 3.3 Å respectively. The G274I mutant exhibited the most compact state, while G275I showed the most expanded state. Structural overlays revealed that the kink in TM6 caused by G275I shifted to a glycine residue above its wild-type position, likely due to the increased bulkiness of the isoleucine substitutions, which induced full bending of the TM6-7 region. These structural changes correlated with hyperpolarizing (G274I) voltage sensitivity shifts and absence of NLC (G275I), as demonstrated previously (*16*). D) Membrane area changes associated with these mutations mirrored those observed during membrane thinning, reinforcing the functional connection between prestin and lipid bilayer mechanics, and the importance of TM6-7 reorganization in prestin’s conformational cycle. E) The anion-binding site remained Cl^-^bound in both mutants, consistent with lipid nanodisc structures, further supporting the conserved role of the anion in the electromotive cycle. F) Structural explanation for the distinct effects for the G274I and G275I mutants. Residue 274 is lipid facing, and the G274I mutation increases the propensity of TM6 helix formation. This increase in helical stability correlates with the compact conformation and the hyperpolarizing shifts in prestin’s NLC response. Residue 275 is facing TM7, and the G275I mutation leads to steric clashes with residue 292, thereby preventing the straightening of TM6. This correlates with the formation of the expanded conformation, and the lack of NLC response.

We then evaluated the dynamic behavior of TM6-7 using hydrogen-deuterium exchange mass spectrometry (HDX-MS) under the same lipidic conditions as before (**fig. 2A**; **fig. S6, Table S1**). Our results supported the observed conformational dynamics and confirmed the impact of the Glycine to Isoleucine mutations on TM6-7 flexibility (**fig. 2A**). Residues 275-292 in the TM6-TM7 linker following the conserved Gly274 and Gly275 hinge exhibited heightened flexibility in nanodiscs, as compared to detergent reconstituted prestin. Moreover, treatment of nanodisc-reconstituted prestin with α-cyclodextrin to induce tension and membrane thinning further increased the flexibility of the TM6 hinge region. G274I faces the lipid bilayer and increases the folding stability of TM6 (**fig. 2, A and F**), facilitating the formation of a “straightened” TM6 and the associated compact state, thereby shifting the V_pk_ of this mutant towards hyperpolarized potentials. G275I points instead towards TM7, packing tightly against L292 in the compact conformation. This packing is disrupted in the G275I mutant, where steric clashes between TM6 and TM7 prevent the formation of the straightened TM6 (**fig. 2, A and F**). Notably, covariance analysis (*49*) revealed evolutionary coupling for amino acids at positions G275 and L292. This coupling is evident in prestin sequence alignments, where one residue typically has a bulky side chain while the other has a small or absent one **(fig. 2B**). We suggest that the altered TM6 flexibility and bending propensity, as well as the disruption of TM6-TM7 interactions, are directly correlated with the functional behaviors of these mutants. Therefore, evolutionarily conserved G274 and G275 residues emerge as critical structural pivots that facilitate prestin’s conformational transitions in response to membrane forces, setting up the coupling of mechanical and electrical cues for fast electromotility essential for mammalian cochlear amplification.

### Anion binding enables prestin’s gate-core mechanoelectrical coupling

Cl⁻ remains bound to prestin’s anion-binding pocket in the expanded conformations induced by either thin bilayers (PC14:1) or the G275I mutation (**Fig. 1, H and I and Fig. 2, D and E**). This observation supports the idea that bound Cl⁻ works like a coupler for gate-core interaction. Further, introduction of a permanent negative charge at the anion binding site (eg. S396D/E and S398D/E) renders prestin insensitive to changes in anionic conditions while S396D restores motor function to near WT levels, as previously proposed (*17, 28, 50, 51*). Whole-cell patch-clamp recordings in HEK293 cells expressing dolphin prestin show that S396D retains a WT-like NLC response with a V_pk_ of −72.5 ± 12.3 mV **(fig. 3A)**. Additionally, the S396D mutant has reduced sensitivity to salicylate, as 10 mM salicylate caused only a slight rightward shift in V_pk_ to −34.5 ± 6.4 mV, unlike the complete NLC abolition observed in the WT (**fig. 3B and fig. S8A**). These results are consistent with the idea that the permanent negative charge introduced by aspartate effectively mimics the coupling role of Cl⁻ on prestin’s electromechanical cycle.

**Fig. 3:**
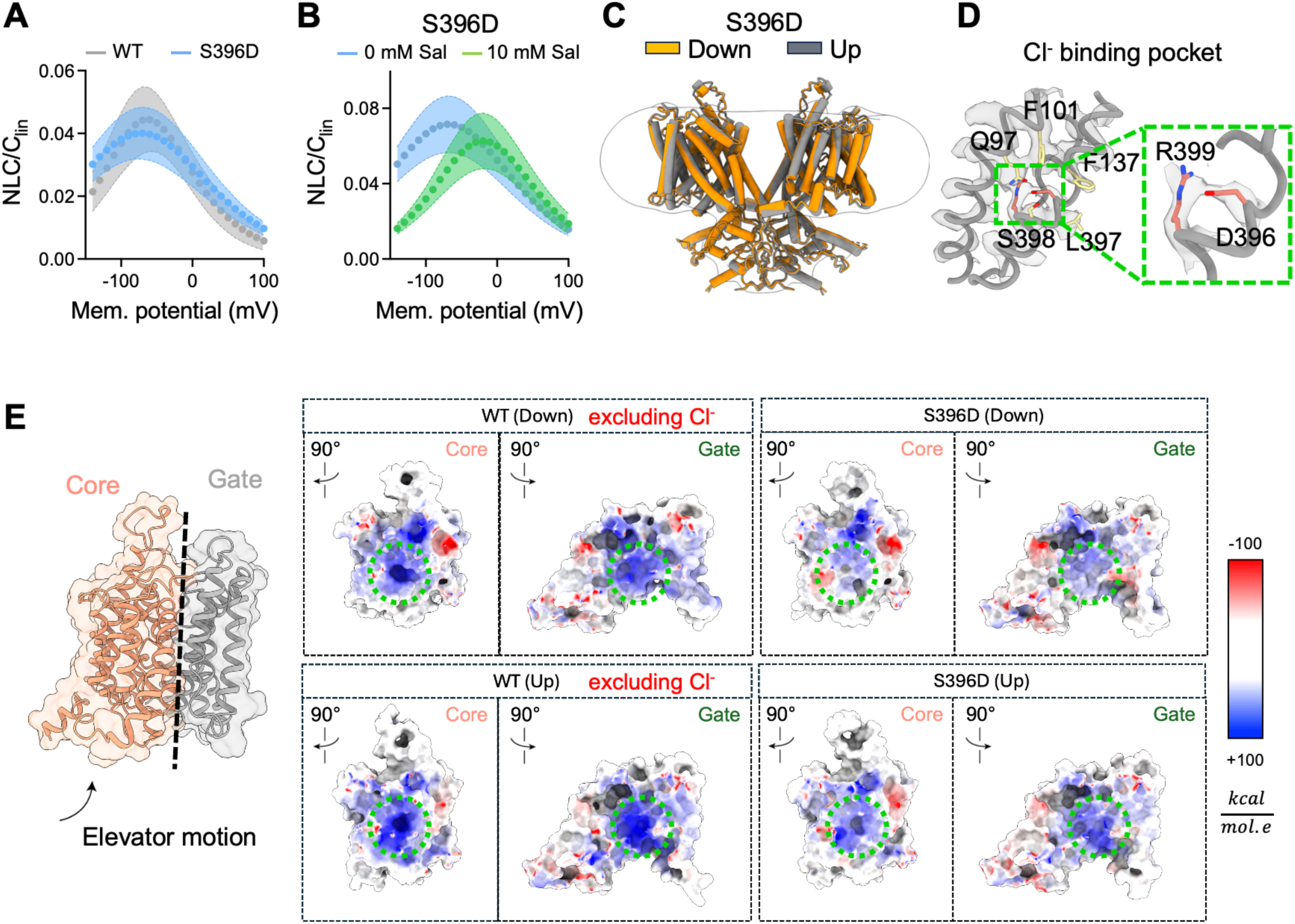
Structural visualization of the role of anions in motor prestin function and electromechanical coupling. A) Whole-cell patch-clamp electrophysiology in HEK293 cells, overexpressing dolphin prestin, demonstrates that replacing a serine with a negatively charged residue, S396D, maintains wild-type like function. The S396D mutant exhibits a V_pk_ of −72 ± 12.3 mV and voltage sensitivity (α) of 0.025 mV^-1^, comparable to wild-type motor Prestin. B) The introduction of the S396D mutation leaves prestin insensitive to salicylate inhibition. C) High-resolution cryo-EM structures of the S396D mutant were solved in detergent and in the presence of Cl^-^ at a nominal resolution of 3.5 Å and 3.6 Å, revealing both Sensor Down and Sensor Up states. D) In the “Up” state, the density of the S396D side chain occupies a position analogous to where Cl⁻ is observed in the wild-type anion-binding site. This demonstrates that a permanent charge, such as aspartate, can functionally replace Cl⁻ at this critical site. E) Changes in the electrostatic potential at the core-gate interface in the S396D mutant. In the absence of an anion, the anion-binding site and the gate domain remain positively charged, in both expanded and contracted conformations. This positive charge is partially neutralized by the presence D396. The electrostatic potential was calculated using CHARMM-GUI and in the presence of the low dielectric environment in the membrane.

Cryo-EM structures of detergent-solubilized, Cl^-^ -bound S396D prestin, revealed a mixed population of elevator Down and Up states at 3.6 Å and 3.5 Å, respectively (**fig. 3, C and D**). The aspartate side chain was well resolved in the Sensor Up state, where it occupies a similar position to Cl⁻ in the WT anion-binding pocket. Additionally, S396D did not produce a significant effect on the gap (*d*) between the TM3 and TM10 helical dipoles when compared to WT prestin structures in Cl^-^, suggesting that the permanent negative charge does not disrupt the architecture of the anion binding site **(fig. S8C).** In the gate domain of S396D, the transition of elevator Up to Down was accompanied by subtle outward tilting of the TM6-7 helices and an intermediate membrane area expansion (**fig. S8B**), suggesting that the main role of Cl⁻ (and surrogate side chains) is one of charge compensation without significantly affecting the overall architecture of the protein.

The anion-induced charge compensation is particularly relevant at the interface between core and gate domains **(fig. 3E)**. To transition between the expanded and compact conformations, the core domain slides against the gate domain, leading to the tight packing of the interface. This becomes evident when considering the α-carbon distance between S398 near the anion binding site, and L448 in the gate domain (Length “L”; **fig. S8D**). Absent Cl⁻, both the core and gate domains remain positively charged, creating an electrostatic bottleneck. In this scenario, the partial charge compensation from either Cl- or S396D is required to overcome the energetic cost of the conformational transitions. This idea is similar to the “charge-compensation” principle underlying the mechanism of other elevator transporters, where substrate binding reduces the electrostatic penalty of the sliding of the charged core domain against the surface of the gate domain (*52–54*). Interestingly, salicylate, due to its increased size, would sterically prevent the sliding of the core domain against the gate domain, inhibiting prestin function.

### Transporter SLC26A5 does not undergo membrane thickness induced area expansion

Consistent with their evolutionary history, only mammalian prestin exerts a voltage-dependent motor function, while nonmammalian orthologs mediate robust anion exchange in heterologous systems (**fig. 4A**) (*5–7*). We have used zebrafish (*Danio rerio*) SLC26A5, a closely related non-electromotile ortholog (54% sequence identity and 84% sequence similarity; **fig. S9**), as a model to examine the molecular basis for the duality in the responses to membrane perturbations in prestin variants, and to investigate the molecular underpinnings underlying the distinct function profiles of different orthologs. Consistent with previous reports, HEK293 cells transfected with zebrafish SLC26A5 produced clear electrogenic currents in the presence of intracellular SO_4_^2-^, indicating robust anion exchange (**fig. 4A**) (*5*). Moreover, zebrafish SLC26A5 generates a shallow NLC response, (α = 0.011 mV^-1^) with a shifted V_pk_ at 74.1±11.1 mV (**fig. 4B**) (*55*), reflecting zebrafish SLC26A5’s primary role as an anion transporter rather than a piezoelectric motor protein.

**Fig. 4:**
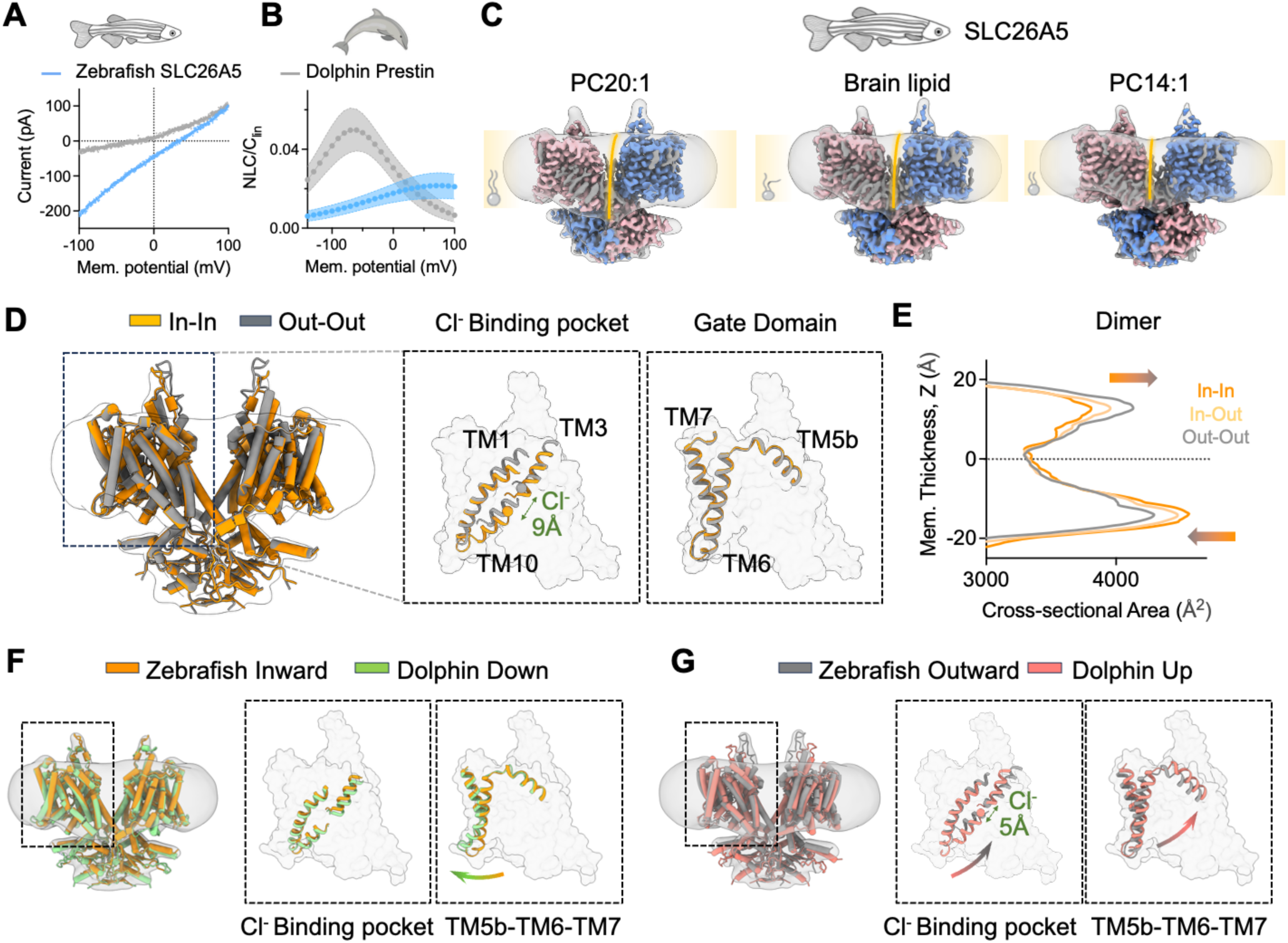
Functional characterization and structural determination of zebrafish SLC26A5. A) Zebrafish SLC26A5 displays robust Cl^-^/SO_4_^2-^ exchange activity, and (B) exhibits reduced nonlinear capacitance (NLC) compared to motor Prestin, functioning primarily as an anion transporter. In HEK293 cells overexpressing zebrafish SLC26A5, the NLC V_pk_ is observed at depolarizing potentials, and voltage sensitivity is reduced (V_pk_ = 74.1±12.1 mV, α = 0.011 mV-1, n = 5 ± SEM). C) Cryo-EM density maps and the overall structure of the zebrafish SLC26A5 homodimer solved in MSP-E3D1 nanodiscs under three lipid conditions: PC20:1 (left), brain lipid extract (middle), and PC14:1 (right). Individual monomers are shown in blue and salmon. Yellow lines indicate membrane thickness. D) Overlay between zebrafish dimers in symmetric in-in and out-out conformations shown as cartoon. While the core-domain, including the Cl^-^ binding pocket undergoes complete elevator transitions, the gate domain remains static and undergoes minimal movement. This produces a membrane footprint that is characterized by redistribution of areal changes across bilayer leaflets (E) and is distinct from the uniform area expansion observed for motor prestin. F) Structural comparison of dolphin prestin in the Sensor Down conformation and zebrafish SLC26A5 in the inward facing conformation. The structures are overall highly similar, with the main difference being the outward bending of TM6 for dolphin prestin only. G) Structural comparison of dolphin prestin in the Sensor Up conformation and zebrafish SLC26A5 in the outward facing conformation. While the Cl^-^-binding site of zebrafish SLC26A5 goes beyond that of dolphin prestin to be exposed extracellularly, the gate domain of zebrafish SLC26A5 acts as a static entity. This contrasts with dolphin prestin, where the gate domain, particularly TM5b-TM6-TM7, reorients itself along with the partial elevator movement of the core domain.

We then compared determined zebrafish SLC26A5 structures in thin lipid nanodiscs (PC14:1), “mixed/regular lipids” (brain lipid extract), and thick lipid nanodiscs (PC20:1) (**fig. 4C, fig. S10 and S11**) to compare the response of zebrafish and dolphin prestin to membrane perturbation at a molecular level. Consistent with the shallow NLC response of zebrafish SLC26A5, no lipid condition displayed a stabilized conformational state for the TMDs, with all samples exhibiting extensive conformational heterogeneity in the elevator domain (**fig. S11).** Specifically, we observed symmetric inward-inward facing, asymmetric inward-outward facing, and symmetric outward-outward facing (for PC14:1 only) dimeric complexes (**fig. 4D**). Furthermore, rather than leading to an area expansion/contraction across both leaflets, as observed for dolphin prestin, the transport producing elevator transitions observed for zebrafish SLC26A5 result in an alternate leaflet-specific redistribution of area changes (**fig. 4E**), similar to previous observations on pendrin (SLC26A4) (*21*).

Given the heterogeneous nature of all zebrafish SLC26A5 structures, we performed an additional round of focused 3D-classification of individual monomers in brain-lipid nanodiscs (**fig. S11B and fig. S12**). This allowed us to resolve an additional intermediate conformation of the TMD characterized by subtle rearrangements of TMs 1,3, and 8 (**fig. S12A**). Importantly, while the inward to intermediate transition produces a subtle contraction that is limited to the intracellular leaflet, the full transition to the outward conformation leads to a substantial expansion in the extracellular leaflet, highlighting a critical difference in the membrane footprint of the conformational cycle for transporter vs motor prestin (**fig. S12B**). Moreover, while area expansion for motor prestin is characterized by concerted motion of both core and gate domains, the transitions of zebrafish SLC26A5 are dominated by Z-direction movements of the core (elevator) domain that leads to an alternately exposed anion-binding site without movements in the gate domain (**fig. S13**).

## Discussion

Prestin, a piezoelectric-like motor protein in OHCs, mediates somatic electromotility in mammals by the reversible conversion of electrical signals into mechanical motion (*10, 14*). This process is pivotal for sound amplification in the cochlea, especially at high frequencies, where OHC-driven electromotility is thought to enhance basilar membrane vibrations. The efficiency of this amplification is contingent upon the rapid response of prestin, operating within the electrical constraints set by the RC time constant of the cell (*56, 57*). Recent advanced measurements *in vivo* and *in vitro* have extended prestin’s functionality due to voltage stimulation to frequencies approaching 19-20 kHz (*58, 59*). At ultrasonic frequencies (>20 kHz), prestin may also contribute to basilar membrane vibrations either passively by modulating the system’s mechanical properties (*60*) or actively if mechanical forces modify its charge movement (“gating charge”) (*61*).

Our data unequivocally show that prestin is inherently mechanosensitive. Membrane thinning drives a clear conformational transition from a compact to an expanded state, with up to ∼10% change in cross sectional area and concomitantly biases the voltage sensor from Sensor-Up to Sensor-Down, respectively. This expansion/contraction in the plane of the bilayer couples membrane forces to transmembrane charge movements, offering a direct route for membrane mechanics to tune prestin’s electromechanical cycle at high stimulus frequencies. Prestin‘s mechanosensitive response mirrors bilayer-coupled gating seen in bona fide mechanosensitive channels (MscL, MscS, Piezo1) (*34, 36, 62, 63*), but has evolved here through natural selection in a transporter-derived scaffold, ultimately repurposed for rapid motor output. Still, comparison between mammalian and non-mammalian (i.e. zebrafish) SLC26A5 underscores the fact that strong bilayer coupling is not a generic feature of the SLC scaffold (**fig. 5A**). Unlike motor prestin, membrane thinning does not produce a uniform net area change across leaflets for zebrafish SLC26A5; rather, it alters area alternatively between leaflets and shifts the state distribution toward outward-facing conformations. Therefore, transporter homologs execute elevator movements with minimal total area change, whereas prestin undergoes coordinated, leaflet expansion/contraction consistent with area-motor behavior, highlighting an evolutionary specialization that enables tight coupling between prestin’s areal expansion and membrane mechanics.

**Fig. 5:**
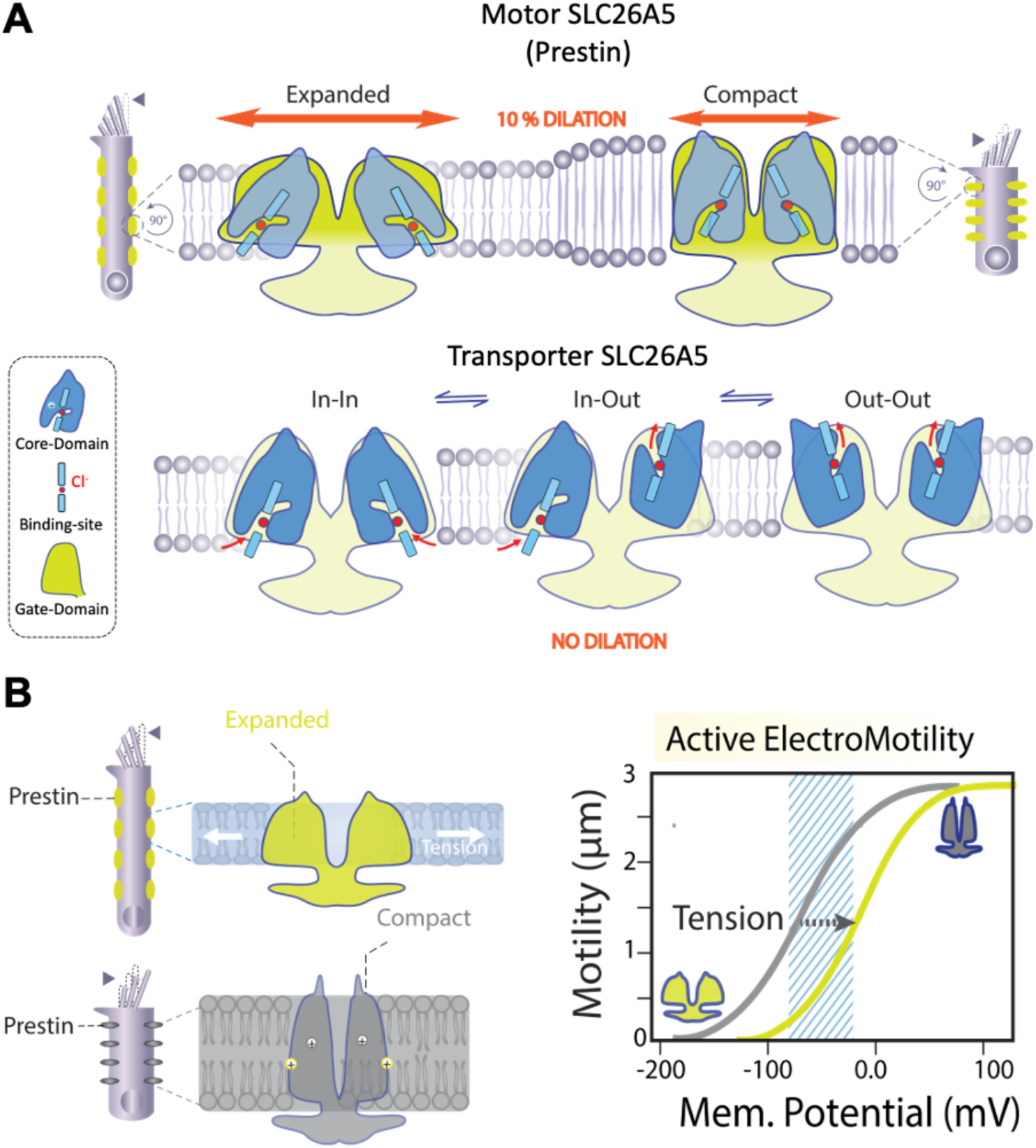
Evolutionary adaptations of motor Prestin for sensing membrane-mediated mechanosensitivity. A) Structural and functional distinctions between Prestin’s electromotility and the SLC26A5 transport cycle. Motor Prestin directly responds to membrane tension and bilayer mechanics by undergoing a uniform change in in-plane area (∼ 10 % dilation), made possible by a flexible gate domain that follows the membrane footprint during tension or thinning. In contrast, the transporter predominantly occupies an asymmetric conformation, with one monomer in an outward-facing state and the other in an inward-facing state. These alternating states, which drive substrate transport, do not produce any net change in membrane area. In the transporter, the gate domain is rigid and permits only an elevator-like sliding motion of the core domain during the transport cycle, preventing the uniform area expansion seen in motor Prestin. Taken together, these observations show that only motor Prestin has evolved a structurally flexible gate domain capable of generating uniform membrane-area changes in response to mechanical cues. B) Our molecular model postulates that motor prestin’s mechanosensitivity allows it to finely tune its electromotility in response to membrane-mediated mechanical forces. As membrane stretch shifts prestin from a compact (Sensor Up) to an expanded (Sensor Down) conformation, gate/core electromechanical coupling provides an adaptive gain control mechanism. This dynamic regulation could prevent saturation at high sound intensities while simultaneously boosting amplification at lower intensities, optimizing sound perception.

Electrostatics at the anion-binding site further tune this cycle. Large anions that occlude the cavity suppress both prestin motor behavior and transporter activity, indicating shared steric control over core–gate coupling. Removing chloride abolishes the NLC response in zebrafish SLC26A5, implicating anion occupancy as a conserved regulatory mode (*64*). Specifically, Cl⁻ acts as a passive cofactor that enables gate-core coupling: a fixed negative charge at S396 (S396D) can substitute for Cl⁻, indicating that charge compensation at this locus is sufficient to support voltage-to-motion coupling. These findings position both electrostatics and bilayer mechanics as joint determinants of prestin’s conformational landscape and can be explained on the basis of a global model of prestin mechanosensitivity (**fig. 5B**): membrane thinning/tension imposes a mechanical bias toward prestin’s expanded state, while anion binding-site electrostatics tune the energy difference between states. Given prestin’s high lateral density at the OHC basolateral membrane and the fact that the bilayer is a shared elastic medium, prestin’s area changes are expected to generate local thickness/tension fields that could bias neighboring motors, providing a plausible mechanism for a bilayer-mediated cooperativity across the OHC membrane.

## Acknowledgments

We thank Drs Omid Bavi, Charles Cox, Boris Martinac, Pancho Bezanilla, and members of the Perozo and Bavi labs for a healthy exchange of ideas and comments on the manuscript. We are grateful to Dr. Peng Shi for sharing the *Tursiops truncatus* Prestin plasmid and Dominik Oliver for the *Danio rerio* SLC26A5 construct. We also thank James Fuller, Joe Austin II, and Tera Lavoie at the University of Chicago Advanced Electron Microscopy Facility and Judy Su and David Strugatsky at California NanoSystems Institute (CNSI) at UCLA for microscope maintenance and training. This work was supported by NIDCD grant R01 DC019833 (to E.P.) and NIH/NIDCD DC017482 (to K.H.). This research was in part supported by the National Cancer Institute’s National Cryo-EM Facility at the Frederick National Laboratory for Cancer Research under contract HSSN261200800001E.

## Author contributions

N.B. and E.P. conceived the project, provided reagents, and with P.H. wrote the manuscript with input from all other authors. N.B. and P.H. expressed and purified the protein, prepared cryo-EM grids, processed the cryo-EM data, built and refined the atomic models, managed cell culture, and, along with K.H., performed and analyzed the electrophysiological experiments. N.B. and P.H. also carried out molecular cloning, mutagenesis, and created all expression constructs with W.M.. N.B., P.H., and M.Z. performed EM data collection. X.L. and T.S. performed and analyzed the HDX-MS. All authors analyzed the data.

## Competing interests

Authors declare no competing interests.

## Materials and Methods

### Cell lines

The cell lines used in this study are similar to the ones used previously. Suspension HEK293S GnTI^-^cells were obtained from ATCC (ATCC CRL-3022) and used for large-scale expression of dolphin prestin and zebrafish SLC26A5. These cells were grown at 37°C and 7.8% CO_2_ in FreeStyle™ 293 expression medium (Gibco Fisher Scientific) supplemented with 10 µg/ml penicillin/streptomycin (Company) and 2% heat-inactivated fetal bovine serum (FBS) (Company). Adherent HEK293T cells were obtained from ATCC (ATCC CRL-3216) and used for patch-clamp electrophysiology experiments. They were cultured at 37°C and 5% CO_2_ in in Dulbecco’s Modified Eagle’s Media (DMEM, Gibco Fisher Scientific) supplemented with 10% heat-inactivated FBS. Suspension Sf9 insect cells (Thermo Fisher, 12659017) were used for baculovirus production. They were grown at 28 °C in SF-900 II SFM (Thermo Fisher, 10902096) medium supplemented with 10% heat-inactivated FBS and 10 µg/ml Gentamicin (Company).

### Generation of prestin constructs

The cloning strategy for dolphin prestin was described previously (*16*). A similar strategy was used for zebrafish SLC26A5, which was a generous gift from the Oliver lab(*5*). Briefly, the DNA sequence encoding dolphin prestin or zebrafish SLC26A5 was subcloned into a pEG BacMam vector with a C-terminal HRV-3C protease site, an eGFP, and an 8x-His tag using restriction digestion. This construct was used for P0 baculovirus production using Cellfectin II (Thermo Fisher, 10362100) and the Bac-to-Bac (invitrogen) method. P0 baculovirus was amplified once to generate P1 virus. The coding sequences for prestin genes were further subcloned into a pEG BacMam vector lacking C-terminal tags. This construct was used for electrophysiological experiments. A construct encoding eGFP alone was generated by subcloning the eGFP coding sequence into a pcDNA3.1 vector using restriction digestion. This construct was used as a transfection marker for electrophysiological experiments. The G274I, G275I, and S396D mutations were introduced into the dolphin prestin constructs using KOD DNA polymerase (EMD Millipore, 71085), and the QuikChange site-directed mutagenesis (Agilent) method. The sequence of each plasmid construct was confirmed by DNA sequencing prior to use.

### Protein expression

The same strategy as previously described (*65*) was used for large-scale protein expression and purification of dolphin prestin and zebrafish SLC26A5. HEK293S GnTI- cells were grown at 37 °C until reaching a confluency of 2.4 to 2.8 million cells/ml. At this confluency, cells were infected with 1:10 v/v P1 baculovirus. After 12-24 hours at 37 °C, the cells were supplemented with 10 mM sodium butyrate and transferred to 30 °C for another 36 to 48 hours (until 60 hours post infection). At this point, the cells were harvested via low-speed centrifugation at 6,000 rpm and 4 °C for 20 minutes, resuspended in phosphate-buffered saline (PBS) at pH 7.4, and spun down again. The resulting pellet was flash frozen in liquid nitrogen and stored at −80 °C until further processing.

### Protein purification and **“**on-column**”** nanodisc reconstitution

The same purification protocol was used for dolphin prestin and zebrafish SLC26A5, and a Cl^-^ based buffer was used as the base solution for all purification steps. This buffer consisted of 150 mM NaCl, 20 mM Tris-HCl, 1 mM EDTA, and 3 mM DTT at pH 7.5, and is referred to as Buffer A. To start the purification, the pellets were thawed in a water bath at room temperature, resuspended in Buffer A supplemented with a tablet of cOmplete™ Protease Inhibitor Cocktail (Roche), 1 μg/ml Aprotinin, 1 μg/ml Leupeptin, 1 μg/ml Pepstatin, 0.2 mM PMSF, and 10 μg/ml DNase. All subsequent steps were performed at 4 °C. After resuspension, the cells were dounce homogenized, and 1% n-dodecyl-β-D-maltopyranoside (DDM; Anatrace) and 0.2% cholesteryl hemisuccinate (CHS, Anatrace) was added for protein extraction while rotating for 90 minutes. The cellular debris was separated via ultracentrifugation at 40,000 rpm for 45 minutes, and the supernatant containing the solubilized protein was collected and mixed with GFP-nanobody coupled CNBR-activated Sepharose beads (GE healthcare) for 90 minutes. The GFP-nanobody coupled resin was prepared as previously described (*16, 66*). After binding, the resin was collected and washed in Buffer A with different detergent compositions: the first wash buffer was supplemented with 0.05% DDM and 0.01% CHS; the second was supplemented with 0.05% DDM, 0.01% CHS, and 0.02% glyco-diosgenin (GDN; Anatrace); the third was supplemented with 0.02% GDN. After the third wash step, the protein was either cleaved with HRV-3C protease (*67*) for 2-4 hours (for detergent-based samples) or subjected to a modified on-column nanodisc reconstitution (*68*).

Three different lipid compositions were used for nanodisc reconstitution. “Brain lipid” consisted of pure porcine brain lipid extract (Avanti Polar Lipids); “Thin lipids” consisted of 90% PC14:1, 1,2-dimyristoleoyl-sn-glycero-3-phosphocholine (Avanti Polar Lipids) and 10% brain lipid extract; “Thick lipids” consisted of 90% PC20:1 1,2-dieicosenoyl-sn-glycero-3-phosphocholine (Avanti Polar Lipids) and 10% brain lipid extract. In all cases, lipids solubilized in chloroform were first dried using rotary evaporation, resuspended in Buffer A, and solubilized using a final concentration of 3% GDN, sonication, and freeze-thaw cycles until the solution was completely transparent. The solubilized lipids were either used immediately, or stored at −80 °C. For all nanodisc samples, MSP1-E3D1 was used as the belt protein and prepared as previously described (*68*). A prestin dimer to MSP to lipid ratio of 1:5:500 was used in all cases. Given that we could not measure the exact concentration of prestin bound to resin, the yield was estimated based on previous experience and multiplied by two to ensure addition of sufficient lipids and MSP. To initiate the reconstitution, lipids were added to the protein bound resin, incubated for 5 minutes, and then purified MSP was added. After 20 minutes of incubation, detergent was removed overnight via three successive rounds of Biobeads (Bio-Rad) addition. The next day, the nanodisc reconstituted protein bound to the resin was washed with 50 ml Buffer A containing no detergent and subsequently cleaved with HRV-3C protease for 2-4 hours.

Both detergent and nanodisc samples were eluted post HRV-3C cleavage and concentrated using a 100 kDa MW cutoff centrifugal filter (Millipore concentrator unit) for size-exclusion chromatography (SEC) using a Superose 6, 10/300 GE column (GE Healthcare). The running buffer for detergent samples consisted of Buffer A supplemented with 0.02% GDN, 1 μg/ml Aprotinin and 1 μg/ml Pepstatin, whereas the running buffer for nanodisc samples consisted of Buffer A supplemented with 1 μg/ml Aprotinin and 1 μg/ml Pepstatin. Peak fractions were concentrated to 2-3 mg/ml using a 100 kDa MW cutoff centrifugal filter and immediately used for cryo-EM grid preparation.

### Cryo-EM sample preparation and data collection

The dolphin prestin brain lipid nanodisc sample was frozen onto Quantifoil 200-mesh 1.2/1.3 Copper grids (Quantifoil), whereas UltrAuFoil 300-mesh 1.2/1.3 Gold grids (Quantifoil) were used for all other samples. In all cases, the grids were glow-discharged at 20W for 30-45s using a Solarus Plasma Cleaner (Gatan). A 3.5 µl volume of purified prestin protein was then applied onto each glow-discharged grid and subsequently plunge-frozen into liquid nitrogen cooled liquid ethane using a Vitrobot Mark IV (Thermo Fisher) at 100% humidity and 22°C with a blot time of 3-5s and blot force 1.

The data for the dolphin prestin G275I mutant was collected at the National Cryo-Electron Microscopy Facility at the National Cancer Institute (NCI) on a Titan Krios (Thermo Fisher) equipped with a K3 camera and GIF energy filter (Gatan) at 81,000x magnification, resulting in a physical pixel size of 1.12Å. Movies were collected using a total dose of 50 e^−^ Å^−2^ divided into 50 fractions. The movies for the remaining samples were collected at the University of Chicago Advanced Electron Microscopy Facility on Titan Krios G3 (Thermo Fisher) equipped with a K3 camera and GIF energy filter (Gatan) at 81,000x magnification, corresponding to a 1.068Å physical pixel size. The data was acquired using a total dose of 60 e^−^ Å^−2^ divided into 52 frames.

### Cryo-EM data analysis

All of the cryo-EM data was analyzed using a combination of Relion (*69, 70*) and cryoSPARC(*71*), and a similar processing pipeline was used for all samples. The raw movies were first subjected to beam-induced motion correction using Relion, and the resulting micrographs were transferred to cryoSPARC for patch-based CTF estimation, micrograph curation, and blob particle picking. The particles were extracted using a box size of 256 Fourier cropped to 128 and subjected to one round of 2D-classification to generate templates for template-based particle picking. These particles were again extracted with a box size of 256 Fourier cropped to 128 for several rounds of 2D-classification. Classes that showed prestin-like features were re-extracted without Fourier cropping to generate multiple ab-initio models using C1 symmetry. In all cases, the ab-initio reconstruction job yielded a single good class and additional “junk” classes, which were used as inputs for subsequent rounds of heterogeneous refinement with C1 symmetry. The particles and map corresponding to the best class were then subjected to local CTF and Non-Uniform refinement (*72*) with either C1 or C2 symmetry applied.

In the case of dolphin prestin, no signs of structural asymmetry were observed, and thus, the C2 symmetric reconstruction was used for downstream processing. First, the particles were transferred back to Relion for Bayesian Polishing (*70, 73*), followed by additional CTF-refinement and Non-Uniform refinement in CryoSPARC. In the case of zebrafish SLC26A5, which exhibited structural asymmetry, the C1 symmetric reconstruction was used for downstream processing. Similarly, particles were exported to Relion for Bayesian Polishing, followed by an additional round of 3D Classification without particle alignment, prior to the final CTF-, and Non-Uniform refinements in CryoSPARC. C2 symmetry was applied for maps with symmetric TMDs (ie. inward-inward and outward-outward), whereas C1 symmetry was used for asymmetric TMDs (ie. inward-outward) during the final refinements.

For zebrafish SLC26A5 in brain lipid nanodiscs, we additionally performed a focused 3D-classification strategy to identify additional conformational states of the TMD. To this end, the particles belonging to the final inward-inward and inward-outward complexes were pooled and subjected to a round of non-uniform refinement. The particles were subsequently symmetry expanded along the C2 axis, a local mask around a single TMD was generated, and the signal outside the mask was subtracted. The subtracted particles were subjected to 3D classification without alignment, followed by local refinement in CryoSPARC.

Sample image processing workflows are provided in Figs. S2, S3, S7, and S11. In all cases, local resolution estimation was performed in cryoSPARC. Moreover, the Relion software used to process cryoEM data was curated by SBGrid (*74*).

### Model building and molecular visualization

The models reported in our previous publication (*16*) corresponding to the “Up” (PDB ID: 7S8X) and “Down I” (PDB ID: 7S9B) conformations were used as initial templates for the dolphin prestin models in this manuscript. In the case of zebrafish SLC26A5, ColabFold (*75*) was used to generate an initial template. The predicted structure was first rigid-body fit to the map of zebrafish SLC26A5 in brain lipid nanodiscs, and residues that lacked clear density were removed manually. The corresponding regions include the N-terminus (residues 1-13), C-terminus (residues 728-739), and IVS (residues 587-641). In all cases, the initial template was first subjected to one round of real-space refinement in Phenix (*76*) with morphing and simulated annealing enabled. The outputs from this job were then manually inspected and iteratively refined using a combination of Phenix real-space refinement, Coot(*77*), and Isolde(*78*). Residues with poor side chain density were modeled as alanines. The surface electrostatics for the core and gate domains were calculated using the PBEQ solver (*79*) in CHARMM-GUI(*80*). UCSF ChimeraX (*81*) was used for molecular visualization and analyses. The evolutionary coupling analysis was performed using the EVcouplings web-server (*49*).

### Patch-clamp electrophysiology in HEK293 cells

Adherent HEK293T cells were used for all heterologous expression experiments. The cells were plated 24 hours before transient transfection. For each transfection, 2.5–3 µg of prestin plasmids and 0.4 µg of eGFP plasmid were mixed with 8 µl of Lipofectamine® 3000 (ThermoFisher Scientific) at a reagent-to-DNA ratio of 2:1 in 125 µl of Opti-MEM (Life Technologies). After 20–24 hours of incubation at 37 °C, the cells were transferred to 30 °C to enhance protein expression. Successfully transfected cells were used for nonlinear capacitance (NLC) measurements after 24–48 hours.

For NLC measurements, we followed the same method described previously(*16*). Briefly, membrane capacitance was measured in the whole-cell configuration using a 500 Hz (for zebrafish SLC26A5) or 1 kHz (for dolphin prestin) sine wave stimulus with a 10 mV amplitude, applied during voltage steps 10 ms after the transient response. Voltage steps ranged from −120 mV to +100 mV, with a holding potential of −70 mV. The admittance (Y(ω)) of the system was determined by spectral analysis, and the DC conductance (g) was derived from the steady-state current preceding the sine wave stimulus. The circuit components, capacitance (C), membrane resistance (Rm), and series resistance (Rs), were calculated accordingly(*82*) and explained in details previously (*16*). For the positive pressure experiments (to create membrane tension) in HEK293 cells, cells overexpressing dolphin prestin were subjected to positive pressure of either 0 or 10 mmHg using a High-Speed Pressure Clamp-1 apparatus (ALA Scientific Instruments, Farmingdale, NY, USA), while simultaneously measuring NLC in whole-cell mode.

For all the experiments done for motor (dolphin) prestin, the internal solution typically contains 130 mM CsCl, 2 mM MgCl₂, 10 mM EGTA, 10 mM HEPES, and 5 mM Na₂ATP, with the pH adjusted to 7.2- 7.4 using CsOH and osmolarity set to 290-300 mOsm, supplemented with sucrose if necessary. The external solution typically contains 140 mM NaCl, 5 mM KCl (or less if minimizing potassium currents), 2 mM CaCl₂, 1-2 mM MgCl₂, 10 mM HEPES, and 10 mM glucose, with the pH adjusted to 7.2-7.4 using NaOH and the osmolarity set to ∼300-310 mOsm, supplemented with sucrose if required. Identical solutions were used for NLC measurements involving zebrafish SLC26A5. For electrogenic Cl^-^/SO_4_^2-^antiport measurements of transporter (zebrafish) SLC26A5, the solutions were almost identical to the ones described previously.^43^ Briefly, the internal solution contained 10 mM CsCl, 10 mM Cs_2_SO_4_, 130 mM potassium L-aspartate, 2 mM MgCl_2_, 10 mM EGTA, and 10 mM HEPES. The pH was adjusted to 7.4 using CsOH. The external solution typically contained 120 mM NaCl, 20 mM TEA-Cl, 2 mM CoCl_2_, 2 mM MgCl_2_, and 10 mM HEPES. The pH was adjusted to 7.4 using NaOH. Command voltage ramps from −100 mV to +100 mV or −130 mV to +130 mV were used to measure electrogenic transport currents. Data for all electrophysiological experiments comes from three independent rounds of transfection.

### Modulation of bilayer thickness

Membrane thickness modulation was achieved by incubation of cells with α-cyclodextrin-PC14:1 (thin lipids) or α-cyclodextrin-PC20:1 (thick lipids) complexes. To generate the complexes, 10 mg of PC14:1 or PC20:1 (Avanti Polar Lipids) dissolved in chloroform was dried using rotary evaporation. The dried lipid film was resuspended in 1 ml of external blocking solution supplemented with 20 mM α-cyclodextrin (Sigma-Aldrich) through bath sonication, generating a stock solution with approximately a 2:1 α-cyclodextrin to phospholipid molar ratio. Of note, complete lipid solubilization was not achieved at this high lipid concentration, and a small amount of precipitate remained despite extensive bath sonication. During recordings, the NLC was first measured prior to modulation of bilayer thickness. Subsequently, external blocking solution with the α-cyclodextrin-PC14:1 or α-cyclodextrin-PC20:1 complex was added to the experimental chamber to reach a final concentration of ∼1 mM α-cyclodextrin in the bath solution, and NLC was measured over several minutes until the giga seal was lost (up to 15 minutes).

### Animals

Wild-type FVB/NJ mice (3-7 months of age) were used to obtain OHCs. The euthanasia procedure was approved by Northwestern University’s Institutional Animal Care and Use Committee.

### NLC recording in OHCs

OHCs were isolated as described previously(*83*). Whole-cell recordings were performed at room temperature using the Axopatch 200B amplifier (Molecular Devices) with a 10 kHz low-pass filter. Recording pipettes pulled from borosilicate glass were filled with a solution containing (mM): 140 CsCl, 2 MgCl_2_, 10 EGTA, and 10 HEPES (pH 7.4). Cells were bathed in an extracellular solution containing (mM): 120 NaCl, 20 TEA-Cl, 2 CoCl_2_, 2 MgCl_2_, 10 HEPES (pH 7.4). Osmolality was adjusted to 309 mOsmol/kg with glucose. The intracellular pressure was controlled manually and monitored by a pressure manometer (PM01D, WPI). The electric current response to a sinusoidal voltage stimulus (2.5 Hz, 120 mV amplitude) superimposed with two higher frequency stimuli (390.6 and 781.2 Hz, 10 mV amplitude) was recorded by jClamp (SciSoft Company). C_m_ was determined by a fast Fourier transform-based admittance analysis (*38*).

### NLC data analysis

Voltage-dependent C_m_ data were analyzed using the following two-state Boltzmann equation (**Eq. 1**):

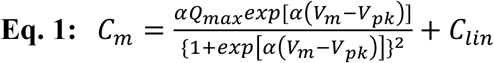

where a is the slope factor of the voltage-dependence of charge transfer, Q_max_ is the maximum charge transfer, V_m_ is the membrane potential, V_pk_ is the voltage at which the maximum charge movement is attained, and C_lin_ is the linear capacitance.

### Hydrogen/Deuterium eXchange Mass Spectrometry (HDX-MS)

Hydrogen-Deuterium Exchange (HDX) combined with Liquid Chromatography-Mass Spectrometry (LC-MS) used purified dolphin Prestin samples. Table 1 provides biochemical and statistical details for HDX in this study per community recommendations (*84*). HDX reactions, quench, injection, and LC-MS were all performed as previously described (*85*). Briefly, HDX was initiated by diluting 2 μL of 20-25 μM prestin stock in an H_2_O buffer into 28 μL of a matching deuterated buffer to reach 93% D-content. Labeling was conducted at room temperature at pD_read_ 7.1. For HDX on nanodisc-embedded prestin in the presence of αCD, the protein was pre-incubated with 10 mM αCD at room temperature for 15 min before dilution into a matching deuterated buffer containing 10 mM αCD to reach 73% D-content.

HDX was quenched at various times, ranging from 10 s to 27 h, by the addition of 30 μL of ice-chilled quench buffer containing 600 mM Glycine, 8 M urea, pH 2.5. For HDX on prestin in nanodisc, the quench buffer also included 3 μL of 0.8% GDN and 3 μL of 300 mg/ml aqueous suspension of ZrO2-coated silica (Sigma-Aldrich, reference no. 55261-U). Quenched reactions were immediately injected into the LC-MS system (Dionex UltiMate-3000 coupled with Thermo Q Exactive mass spectrometer). Protein digestion was performed online by a pepsin/FPXIII mixed protease column prepared in-house. Peptides were assigned by SearchGUI version 4.0.25, and HDX data was processed using HDExaminer 3.1 (Trajan) followed by manual curation.

Changes in free energy of unfolding (ΔΔG’s) were calculated from the area between two deuterium uptake curves after correction for back-exchange (*86*). Residue-level ΔΔG’s were determined from a weighted average of peptide-level ΔΔG’s, excluding the first two N-terminal residues due to rapid back-exchange. We note that, while most peptides exhibited near complete exchange within our labeling times, ΔΔG presented here is considered semi-quantitative, as precise quantification requires a broader and denser sampling of labeling times. Nonetheless, this approach effectively captures relative ΔΔG associated with changes in lipid environment or mutagenesis, and our major conclusions remain independent of this fitting method. The deuteration differences shown in **fig. S7** were adjusted for D-content but not for back-exchange.

### Statistical analysis

Statistical significance (criteria, \**P* < 0.05; \*\**P*<0.005) between different electrophysiological conditions was determined using unpaired or paired Student’s t-tests. R was used for all statistical analysis, and ggplot2 (*87*) was used for plotting.

## Supplementary Figures

**Fig. S1:**
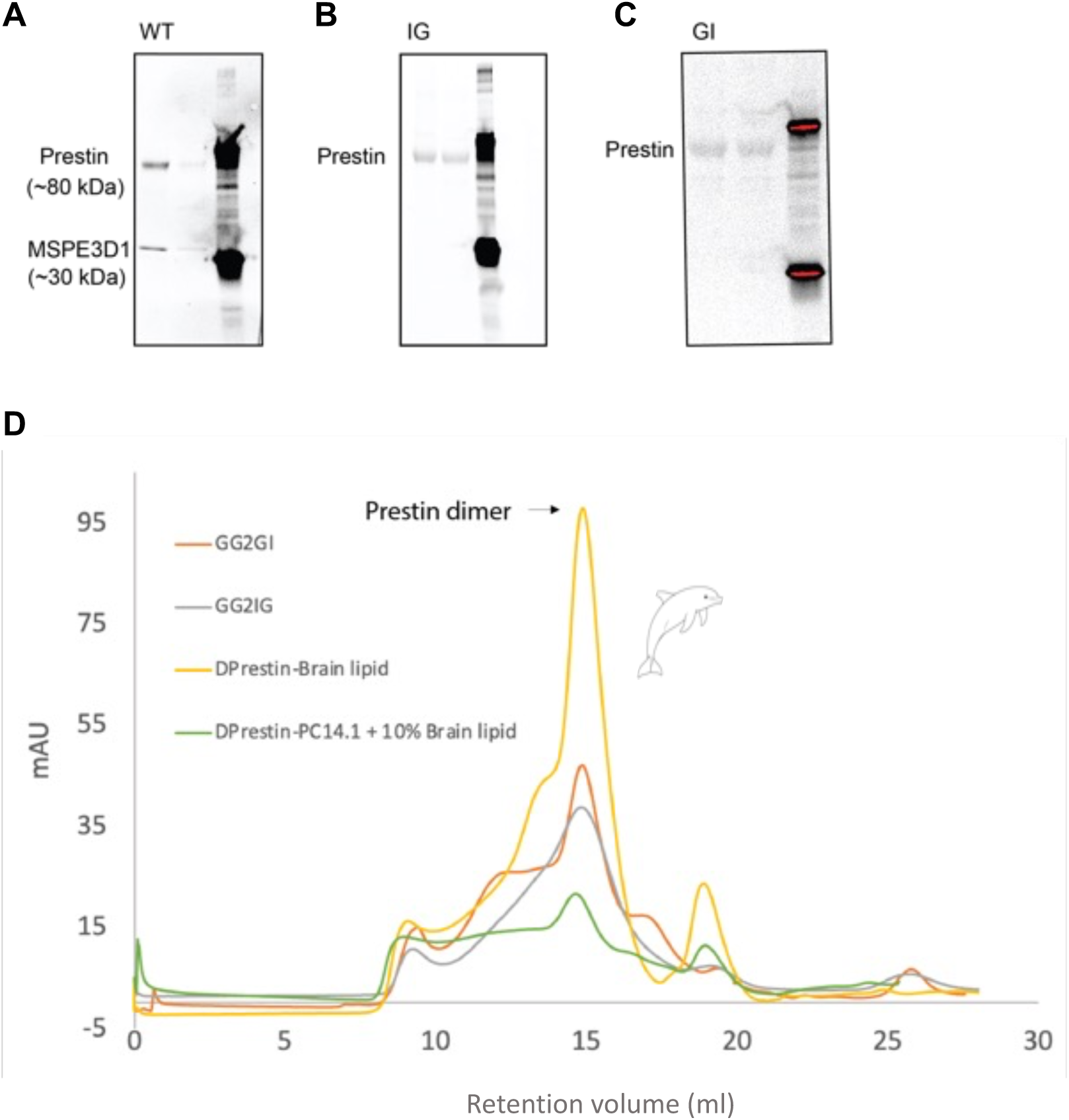
SDS-PAGE and size-exclusion chromatography (SEC) HPLC of purified dolphin prestin mutant in detergent and nanodiscs. A-C) Only representative purification samples are shown for clarity. Prestin monomer elutes at ∼80 kDa (∼160 kDa as a dimer). D) Purification in detergent (A,B) and nanodiscs (C) results in a monodisperse peak using a Superose 6 column, which was subsequently collected and concentrated for sample freezing.

**Fig. S2:**
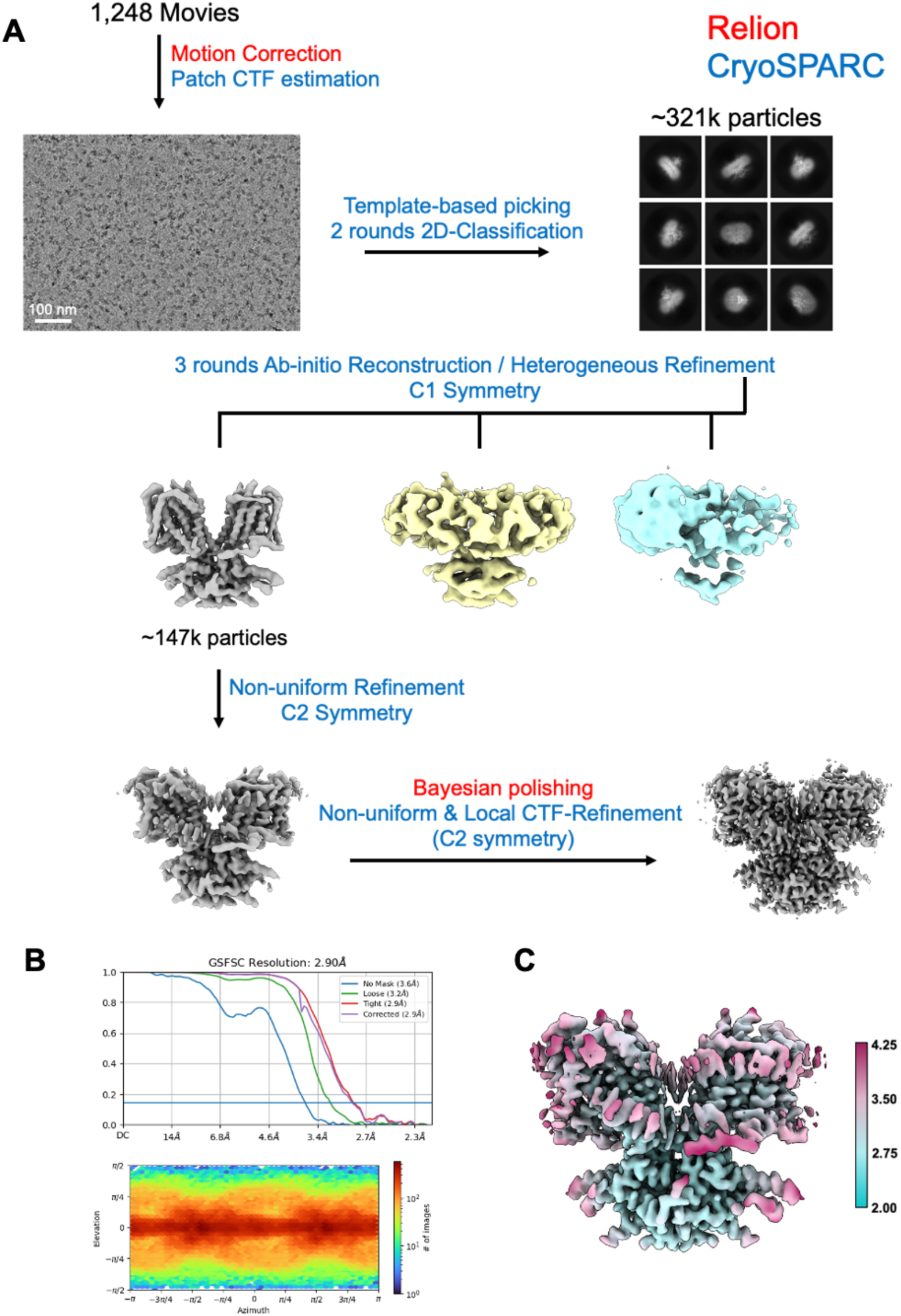
Cryo-EM workflow for dolphin prestin in brain-lipid extract and MSPE3D1 nanodiscs (See Methods). A) Data processing was performed using Relion and CryoSPARC. Motion correction and Patch CTF estimation were followed by template-based particle picking and two rounds of 2D classification. Three rounds of ab-initio reconstruction and heterogeneous refinement were performed with C1 symmetry, followed by non-uniform refinement with C2 symmetry. Bayesian polishing and local CTF refinement further improved map quality, yielding a final 2.90 Å resolution reconstruction, as shown in the FSC and angular distribution plots (B). The local resolution estimates for the final map are illustrated in (C).

**Fig. S3:**
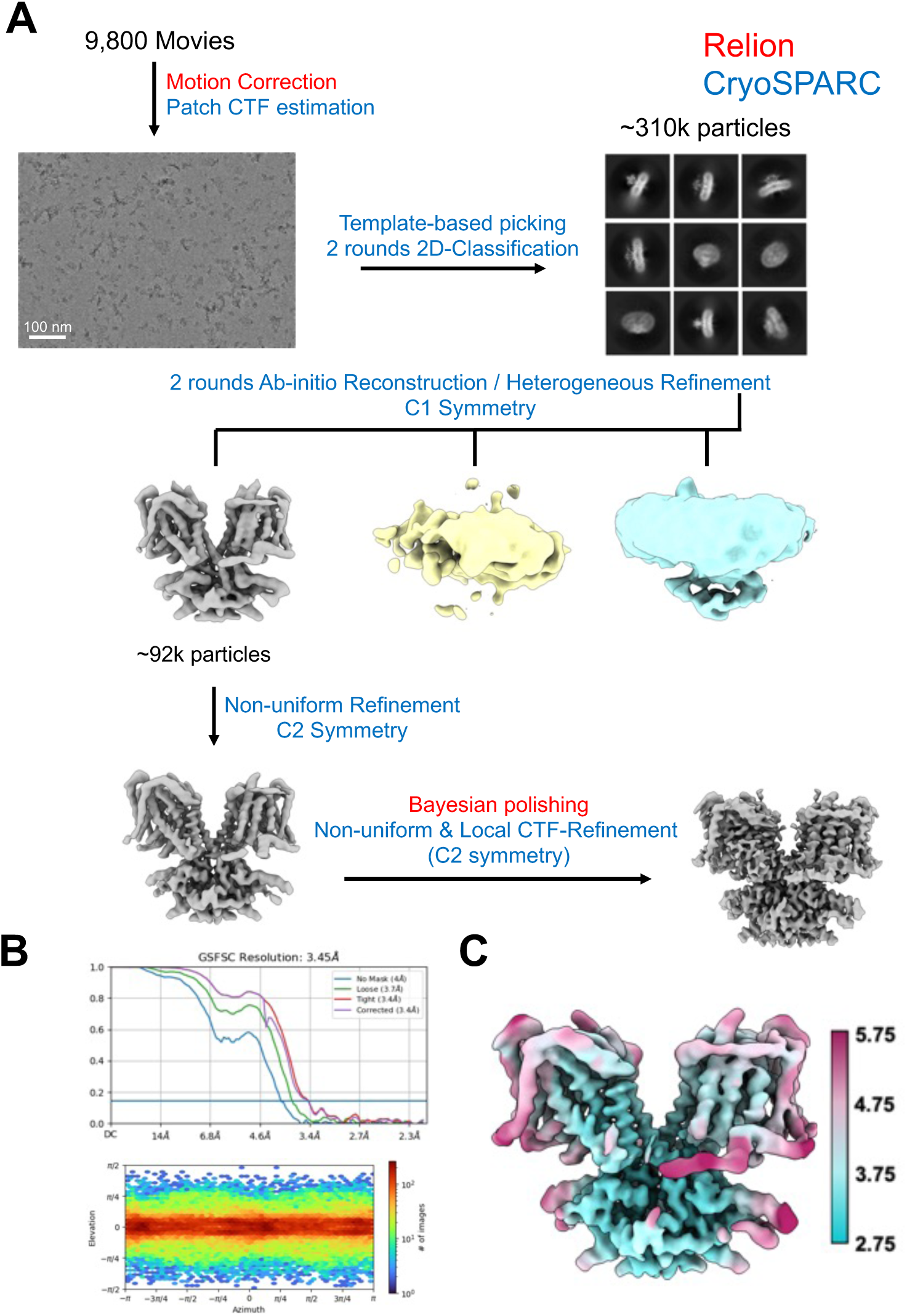
Cryo-EM workflow for dolphin prestin in PC14:1 lipids and MSPE3D1 nanodiscs. Data processing was conducted using Relion and CryoSPARC. The workflow included motion correction and Patch CTF estimation, followed by template-based particle picking and two rounds of 2D classification. Two rounds of ab-initio reconstruction and heterogeneous refinement were performed under C1 symmetry, succeeded by non-uniform refinement with C2 symmetry. Bayesian polishing and local CTF refinement were applied to enhance map quality, resulting in a final reconstruction at 3.45 Å resolution. The FSC and angular distribution plots illustrate the achieved resolution and particle orientation (B). The local resolution estimates for the final map are illustrated in (C).

**Fig. S4:**
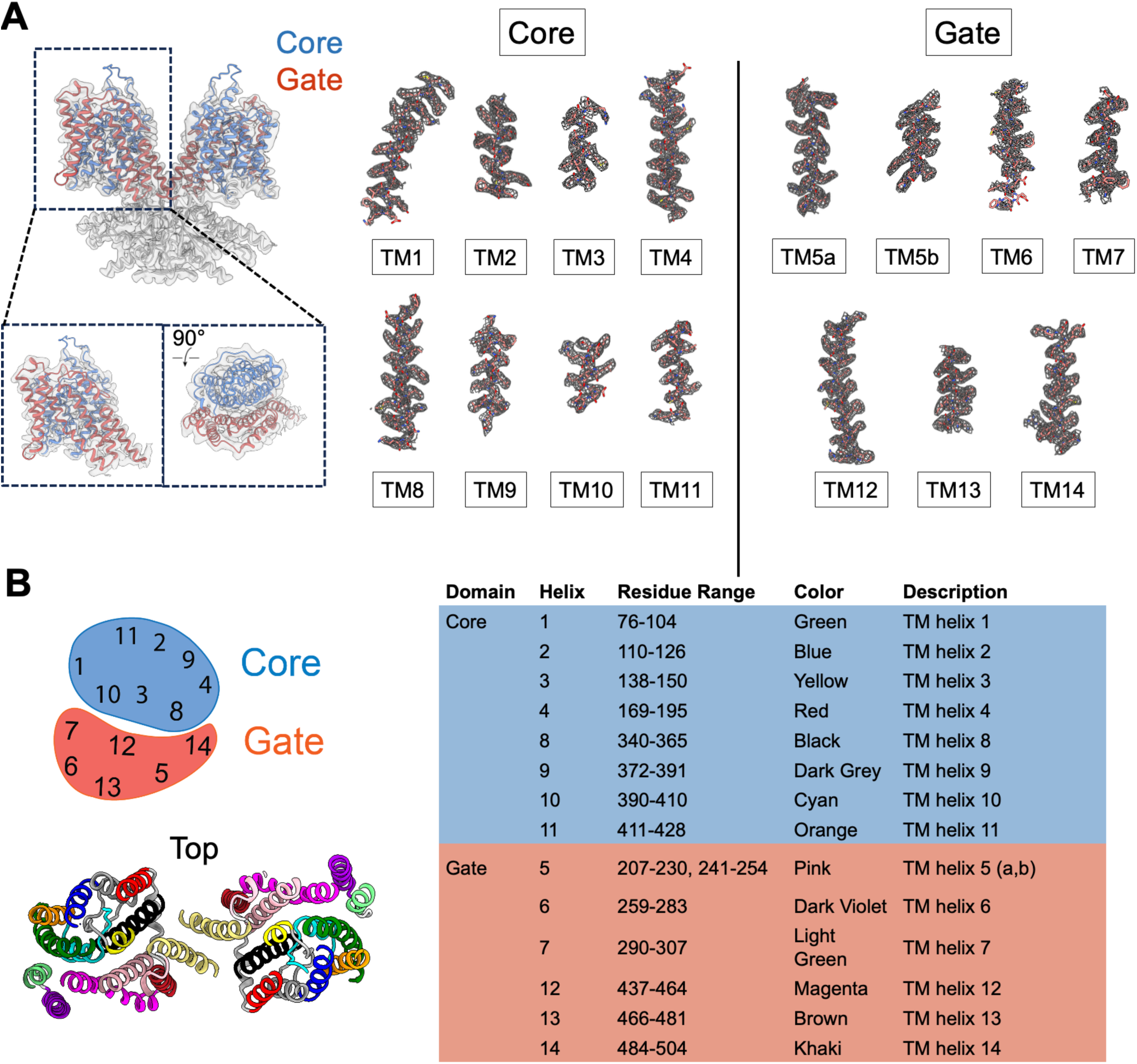
Model-to-density fit and architecture of dolphin prestin as a homodimer. Dolphin prestin in brain lipid extract nanodiscs is shown as an example. A) Density fit for the transmembrane region of dolphin prestin in brain lipid nanodiscs. B) Residue mapping for TM helices (TM1-TM14) in the core and gate domains of prestin.

**Fig. S5:**
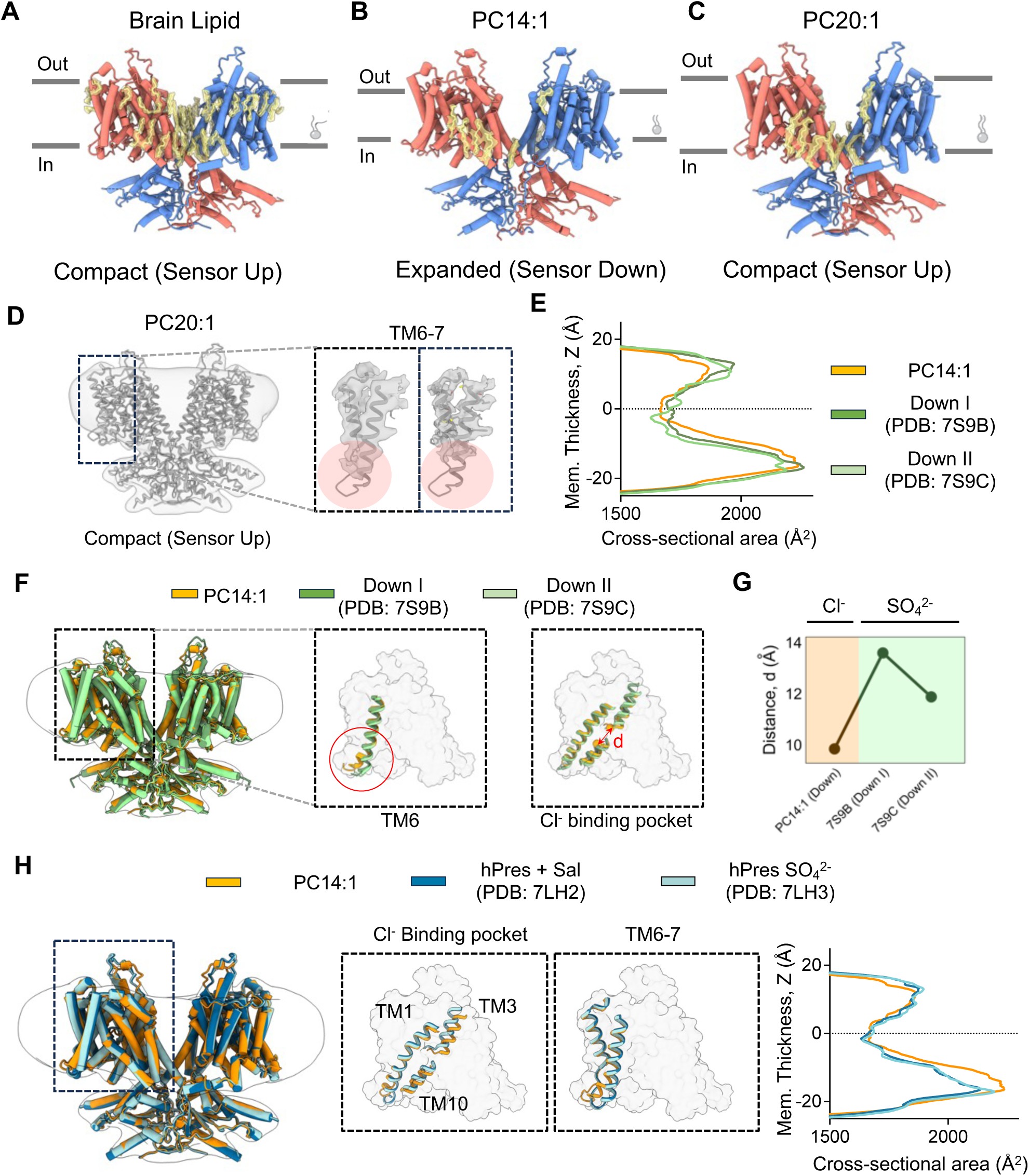
Models of structural lipids based on density maps for dolphin prestin in nanodiscs with various compositions, including (A) brain lipid extract, (B) PC14:1, and (C) PC20:1 (see Methods for detailed composition). Several lipid-like densities were observed at the periphery of prestin’s TMD in the brain lipid nanodisc, indicating its intimate interactions with the surrounding bilayer. The putative lipids were modeled as hydrocarbon chains of varying lengths. For instance, in PC14:1 nanodiscs, putative lipids were modeled as C14 hydrocarbon chains, only into densities of approximately the proper length, whereas in PC20:1 nanodiscs, they were modeled accordingly. D) Close-up view of the cryo-EM density around TM6-7 in the PC20:1 nanodisc sample. The lack of density at the TM6-7 linker suggests substantial conformational flexibility for this region. E) Area comparison for the PC14:1 structure and previously published dolphin prestin structures in the down conformation resolved in the presence of SO_4_^2-^ and detergent (PDB IDs: 7S9B and 7S9C). F) Overlay of the novel expanded structure determined in PC14:1 nanodiscs with previous structures determined in the presence of SO_4_^2-^. Comparison between the structures shows differences in the TM6 bending behavior, and the architecture of the anion binding site. G) Comparison of the dimensions of the anion binding site for different structures in the sensor-down conformation, as measured by the distance (d) between the alpha carbons of residues R399 and F137. The differences in d suggest that the detailed architecture of the anion binding site is defined by the identity of the anion. H) Structural overlay and area comparison of the structure in PC14:1 nanodiscs with previously reported structures of human prestin in the presence of SO_4_^2-^ (PDB ID: 7LH3) and salicylate (PDB ID: 7LH2) in detergent micelles.

**Fig. S6:**
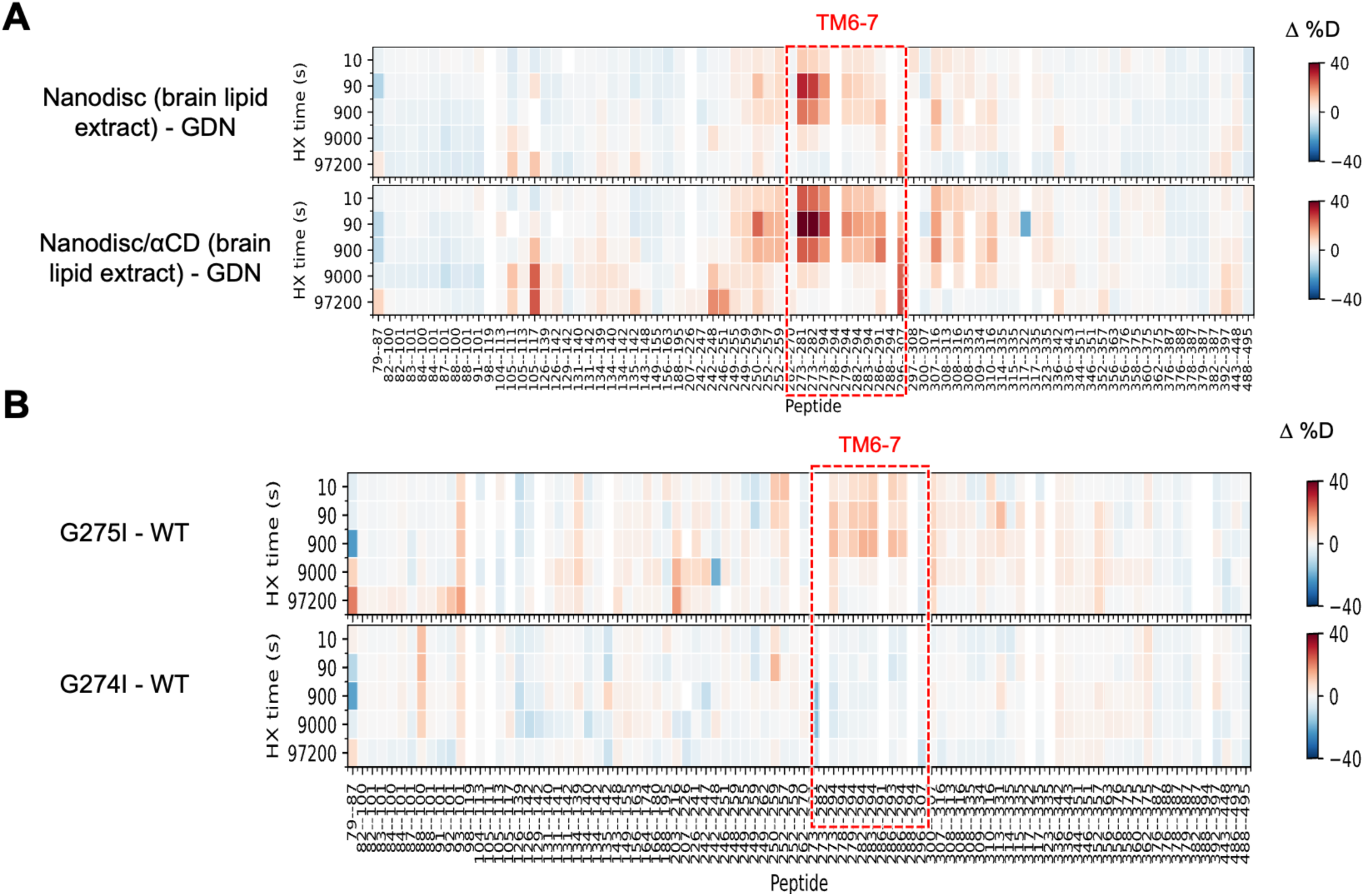
Hydrogen–deuterium exchange mass spectrometry (HDX-MS) of wild-type Prestin in different lipid environments and of prestin mutants in Cl^-^. A) HDX-MS data shows that TM6-7 region dynamics in response to changes in the lipid environment. Heatmaps show differences in deuteration levels at each labeling time for all TMD peptides measured in one of the following conditions compared to wild-type GDN-solubilized prestin: WT prestin in porcine brain total lipid nanodisc with (top) and without (bottom) 10 mM αCD, D) GDN-solubilized G275I (top) and G274I (bottom) mutant.

**Table S1:** Biochemical and statistical details for HDX-MS.

| Dataset | WT, GDN | WT, nanodisc<br>(porcine brain<br>total lipid<br>extract) | WT, nanodisc<br>(porcine brain<br>total lipid<br>extract) + αCD | WT, nanodisc<br>(PC20/porcine<br>brain total lipid<br>extract) | WT, nanodisc<br>(PC20/porcine<br>brain total lipid<br>extract) + αCD | G274I, GDN | G275I, GDN |
| --- | --- | --- | --- | --- | --- | --- | --- |
| HDX reaction<br>details | 20 mM Tris, 360<br>mM NaCl, 3 mM<br>DTT, 1 mM EDTA,<br>0.02% GDN.<br>pD<sub>read</sub> 7.1, 25 °C | 20 mM Tris, 150<br>mM NaCl. pD<sub>read</sub><br>7.1, 25 °C | 20 mM Tris, 150<br>mM NaCl, 10 mM<br>αCD. pD<sub>read</sub> 7.1,<br>25 °C | 20 mM Tris, 150<br>mM NaCl. pD<sub>read</sub><br>7.1, 25 °C | 20 mM Tris, 150<br>mM NaCl, 10 mM<br>αCD. pD<sub>read</sub> 7.1,<br>25 °C | 20 mM Tris, 360<br>mM NaCl, 3 mM<br>DTT, 1 mM EDTA,<br>0.02% GDN.<br>pD<sub>read</sub> 7.1, 25 °C | 20 mM Tris, 360<br>mM NaCl, 3 mM<br>DTT, 1 mM EDTA,<br>0.02% GDN.<br>pD<sub>read</sub> 7.1, 25 °C |
| HDX time course<br>(*: replicated) | 10s*, 30s, 90s*,<br>5min, 15min*,<br>45min, 150min*,<br>27h* | 10s, 90s, 15min,<br>150min, 27h | 10s, 90s, 15min,<br>150min, 27h | 10s, 90s, 15min,<br>150min, 27h | 10s, 90s, 15min,<br>150min, 27h | 10s*, 90s*, 5min,<br>15min*, 150min*,<br>27h* | 10s*, 90s*,<br>15min*, 150min*,<br>27h* |
| HDX control<br>samples | Non-deuterated control; in-exchange control; maximally labeled control |  |  |  |  |  |  |
| In- and back-<br>exchange,<br>mean/IQR (TMD<br>peptides only) | In-exchange: 2.5% / 2.2%; back-exchange: 25% / 13% |  |  |  |  |  |  |
| No. of peptides<br>(TMD only) | 92 | 74 | 74 | 72 | 71 | 84 | 79 |
| Sequence<br>coverage (TMD<br>only) | 75% | 59% | 59% | 63% | 63% | 73% | 73% |
| Average peptide<br>length/redundan-<br>cy (TMD peptides<br>only) | 11.7/2.6 | 11.5/2.1 | 11.5/2.1 | 11.7/2.0 | 11.7/2.0 | 11.9/2.4 | 12.1/2.3 |
| Replicates | 2 (Biological) | 1 | 1 | 1 | 1 | 2 (Technical) | 2 (Technical) |
| Repeatability (SD<br>of the $\Delta\%D/\Delta\#D$<br>between<br>duplicates) (TMD<br>peptides only) | 0.69%/ 0.06 Da | N/A | N/A | N/A | N/A | 1.77%/0.16 Da | 1.99%/0.17 Da |

**Fig. S7:**
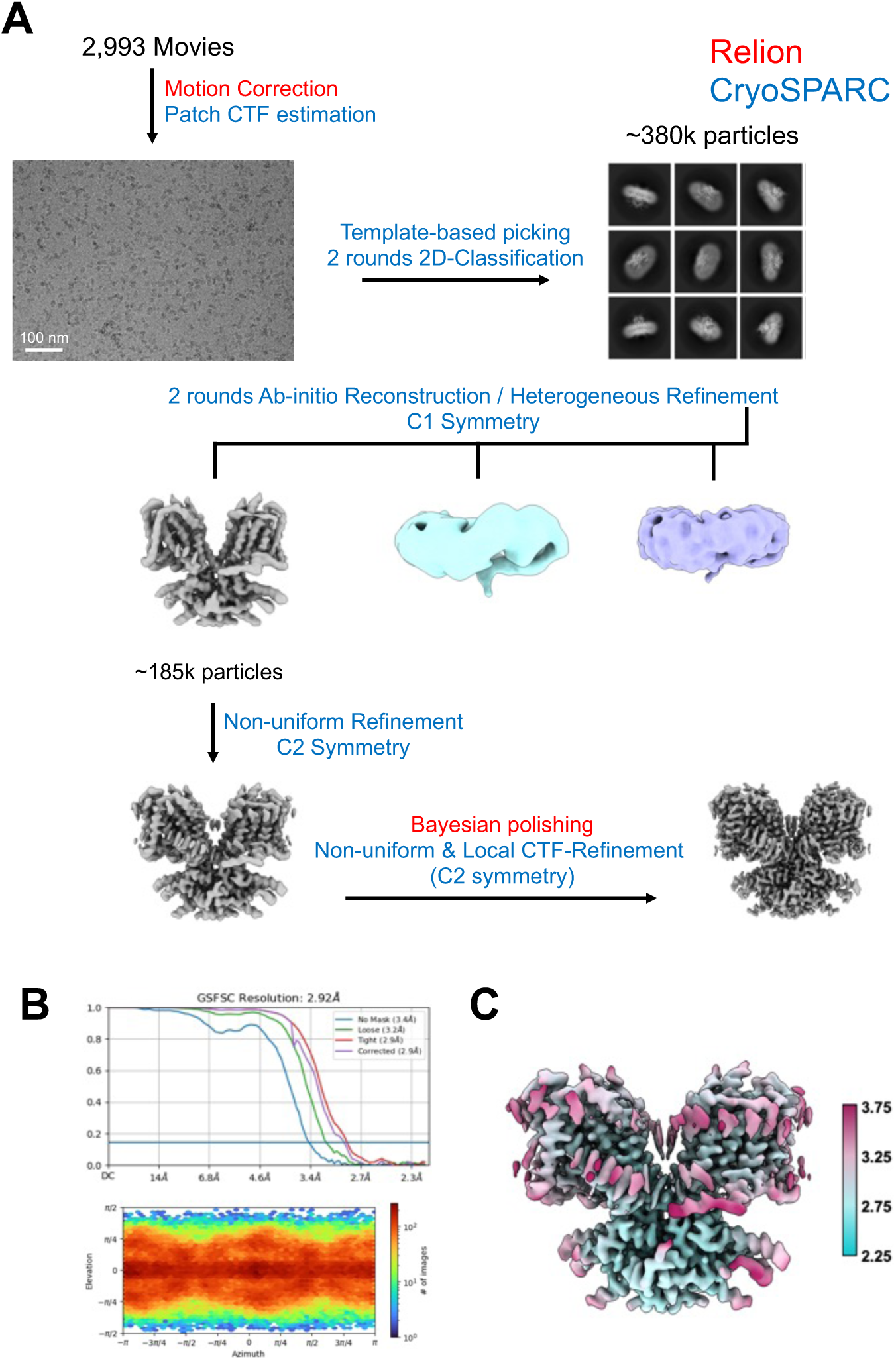
Cryo-EM workflow for dolphin prestin G274I in detergent (GDN). **a)** Data processing was conducted using Relion and CryoSPARC. The workflow included motion correction and Patch CTF estimation, followed by template-based particle picking and two rounds of 2D classification. Two rounds of ab-initio reconstruction and heterogeneous refinement were performed under C1 symmetry, succeeded by non-uniform refinement with C2 symmetry. Bayesian polishing and local CTF refinement were applied to enhance map quality, (**b**) resulting in a final reconstruction at 2.92 Å resolution. The FSC and angular distribution plots illustrate the achieved nominal resolution and particle orientation. **c)** Local resolution estimation for the final reconstruction.

**Fig. S8:**
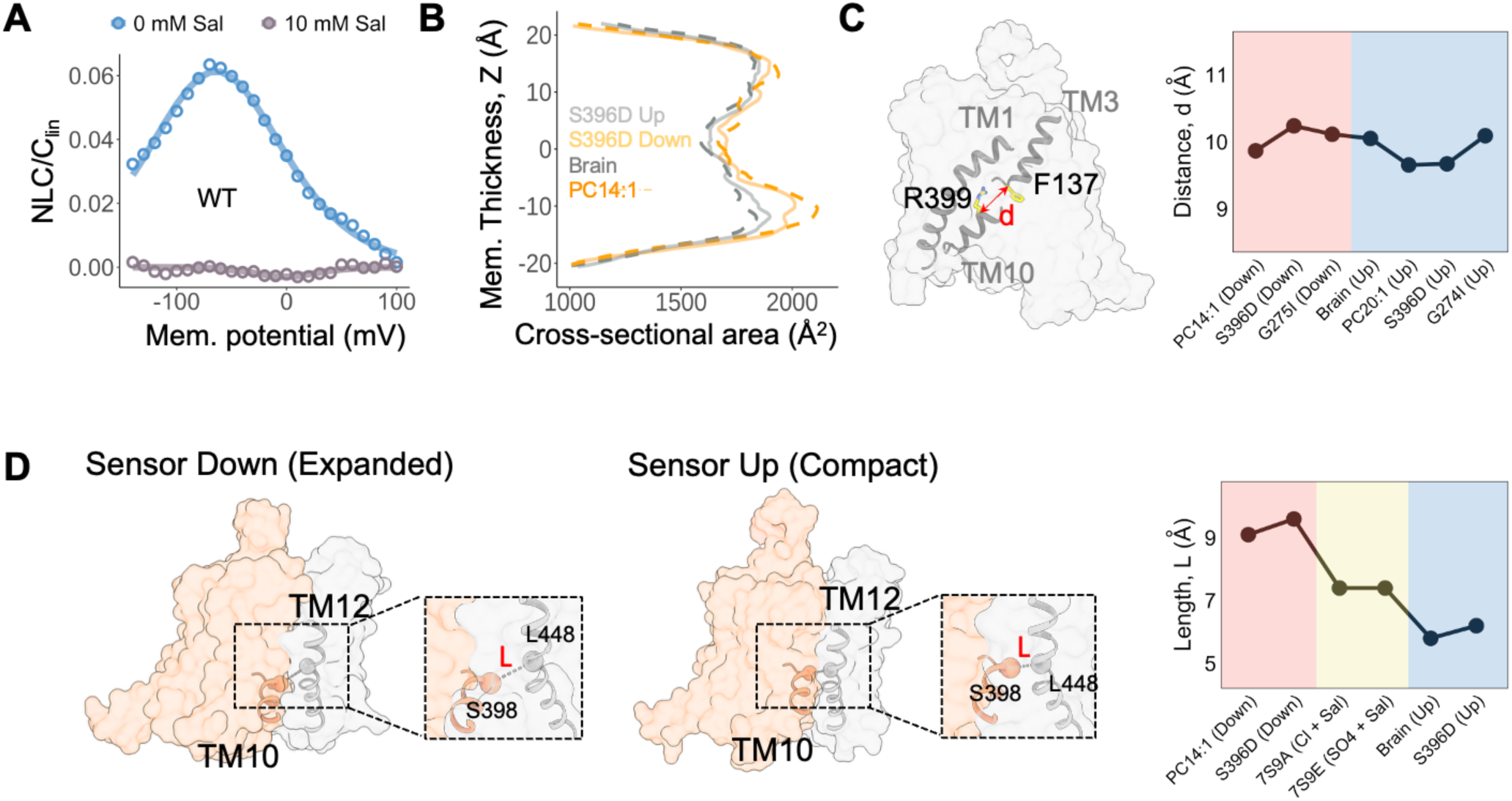
Dynamics of anion binding site size in different states and when occluded with Cl⁻ versus salicylate. A) Whole-cell patch-clamp electrophysiology in HEK293 cells overexpressing dolphin prestin demonstrates that WT prestin is readily inhibited by 10 mM salicylate applied extracellularly. B) Comparison of the intra-membrane cross-sectional area shows that the S396D mutant displays intermediate area expansion. C) *d* represents the distance between the alpha carbon of R399 and F137, which approximately corresponds to the diameter of the anion binding site. The changes in *d* are <1 Å, suggesting that the anion binding site is undisrupted by different mutations. D) Proposed mechanism for the functional effect of partial charge neutralization at the binding site. In order to transition from expanded to contracted conformations, the core and gate domain must slide and pack tightly against each other, as evidenced by changes in length “L”, which represents the distance between the alpha carbon of S398 on the core domain, and L448 in the gate domain. These reorientations are electrostatically unfavorable in the absence of a negative charge, rendering prestin nonfunctional. Moreover, salicylate, due to its increased size, prevents the sliding of the core domain against the gate domain, similarly inhibiting prestin function.

**Fig. S9:**
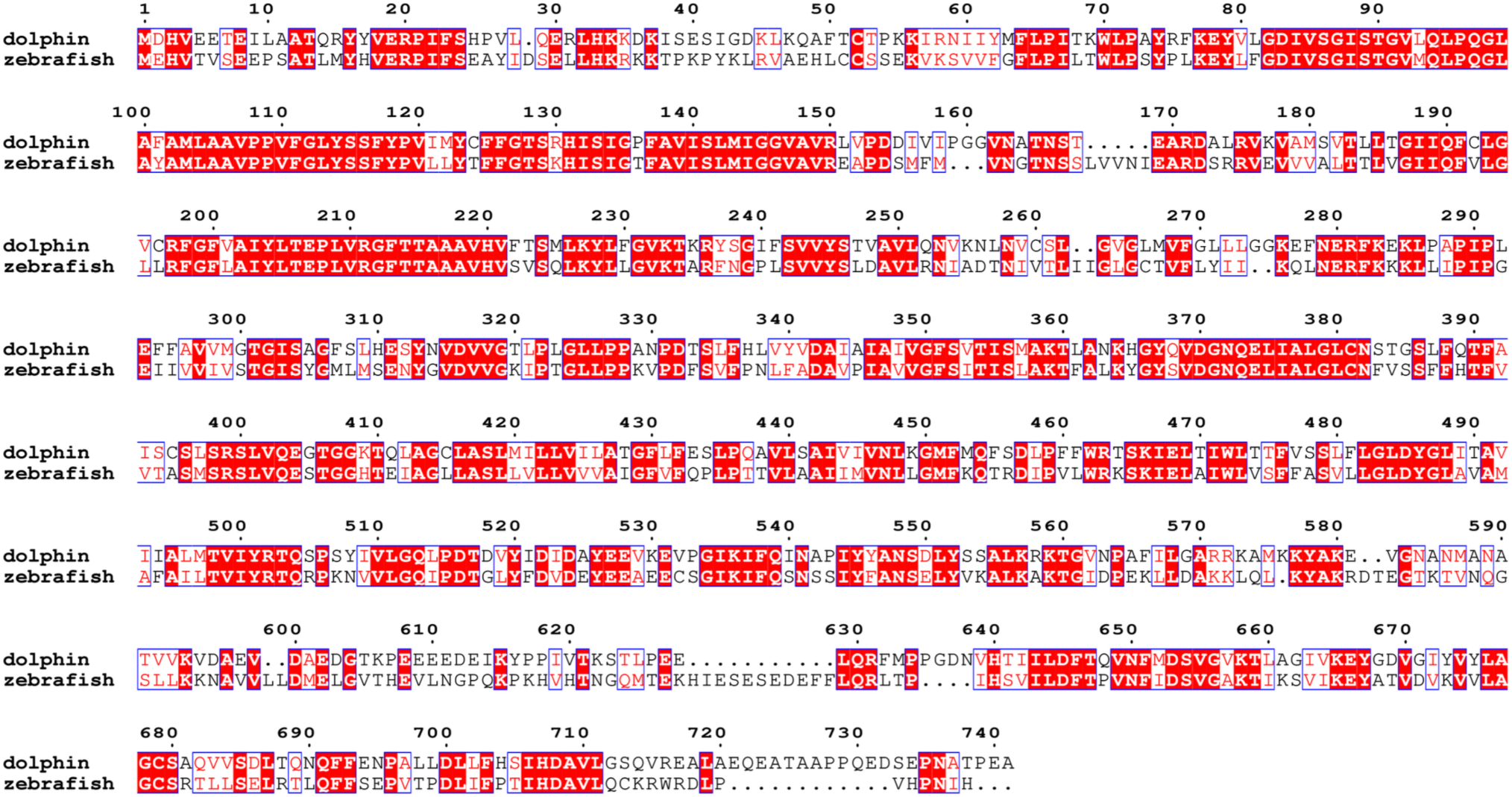
Amino acid sequence alignment for dolphin prestin and zebrafish SLC26A5. Identical residues are highlighted by a red background, whereas similar residues are depicted by red font. Sequence stretches with common physicochemical properties are boxed in blue. The full-length sequences were aligned using MAFFT (*88*), and the alignment figure was generated using ESPript3.2 (*89*).

**Fig. S10:**
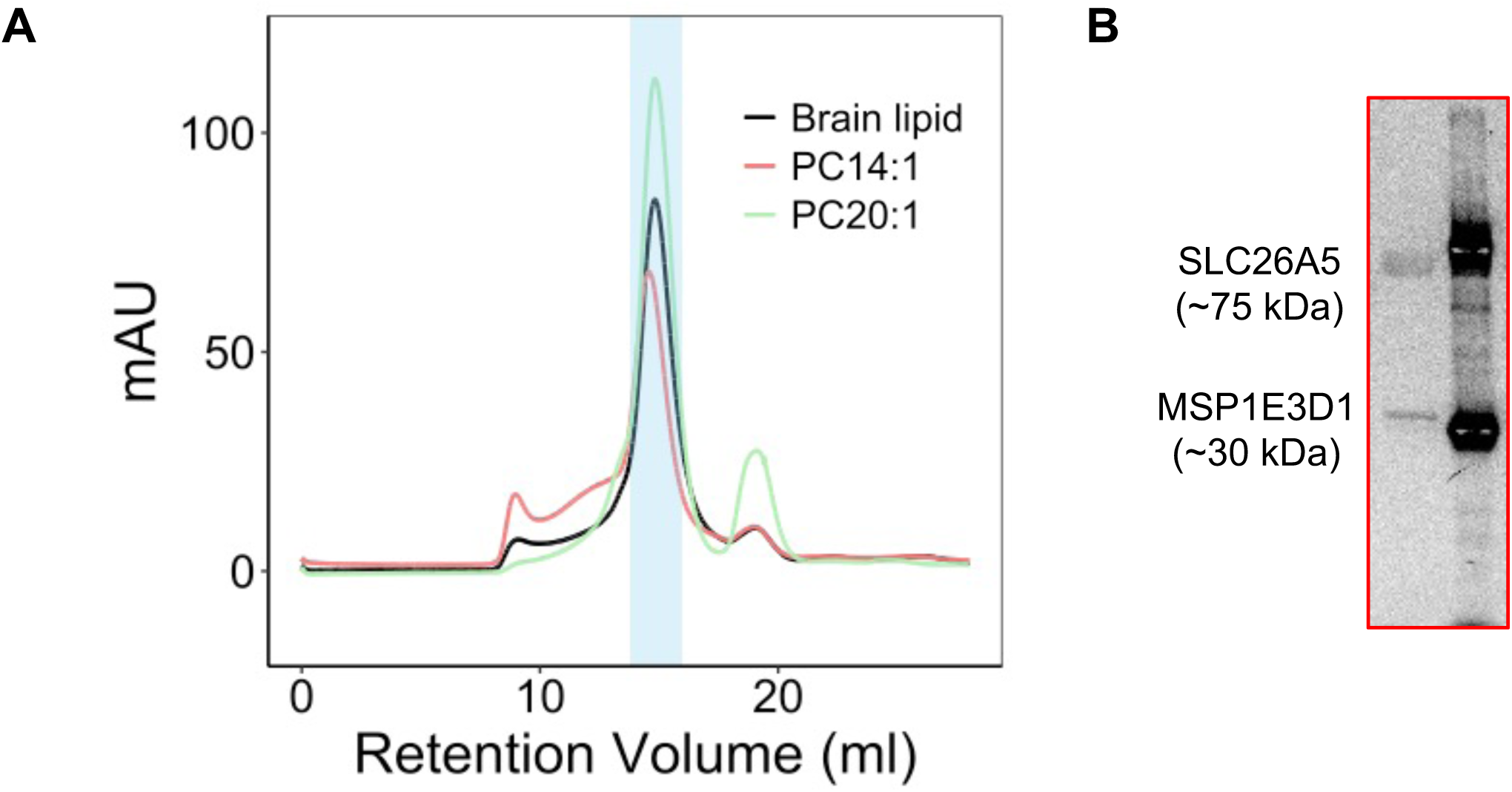
Biochemistry of zebrafish SLC26A5. **a**) A) Size-exclusion chromatography (SEC) HPLC of purified zebrafish SLC26A5 in PC14:1, brain-lipid and PC20:1 lipid composition nanodiscs (MSPE3D1), resulting in a monodisperse peak using a Superose 6 column. The peak was subsequently collected and concentrated for sample freezing. B) Only representative purification samples are shown for clarity. zebrafish SLC26A5 monomer elutes at ∼80 kDa (∼160 kDa as a dimer), while MSPE3D1 elutes at ∼30 kDa.

**Fig. S11:**
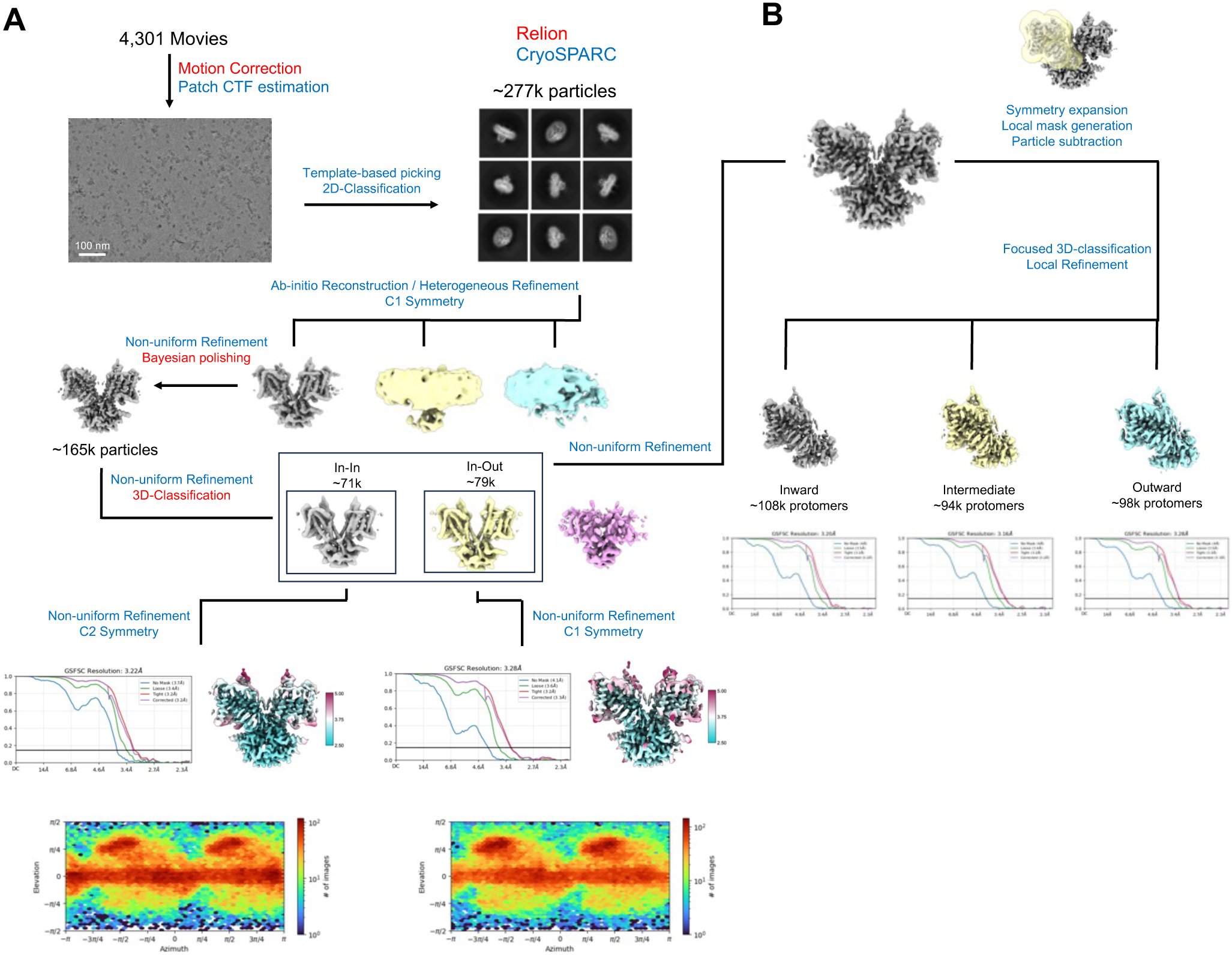
Cryo-EM workflow for zebrafish SLC26A5 in brain lipid nanodiscs. A) Data processing was conducted using Relion and CryoSPARC. The workflow included motion correction and Patch CTF estimation, followed by template-based particle picking and two rounds of 2D classification. Two rounds of ab-initio reconstruction and heterogeneous refinement were performed under C1 symmetry, succeeded by non-uniform refinement with C2 symmetry. Bayesian polishing and local CTF refinement were applied to enhance map quality. After Bayesian polishing, particles were reimported into CryoSPARC for one round of Non-Uniform Refinement and subjected to additional 3D-classification in Relion. The particles corresponding to the inward-inward dimer were subjected to Non-Uniform Refinement using C2 symmetry, producing a map at a nominal resolution of 3.2Å. The particles corresponding to the inward-outward dimer were subjected to Non-Uniform Refinement without symmetry applied, resulting in a reconstruction at 3.3Å nominal resolution. The FSC curves, orientation distribution, and local resolution estimates are shown for both samples. B) Workflow for the additional focused classification of individual protomers. The particles from the inward-inward and inward-outward structures were pooled and subjected to one round of Non-Uniform Refinement. After refinement, particles were symmetry expanded along the C2 symmetry axis, a local mask was generated, and signal outside of the mask was subtracted. The subtracted particles were subjected to 3D-Classification without alignment, and subsequent local refinement produced three distinct maps at 3.2 (inward), 3.2 (intermediate), and 3.3Å (outward) nominal resolution.

**Fig. S12:**
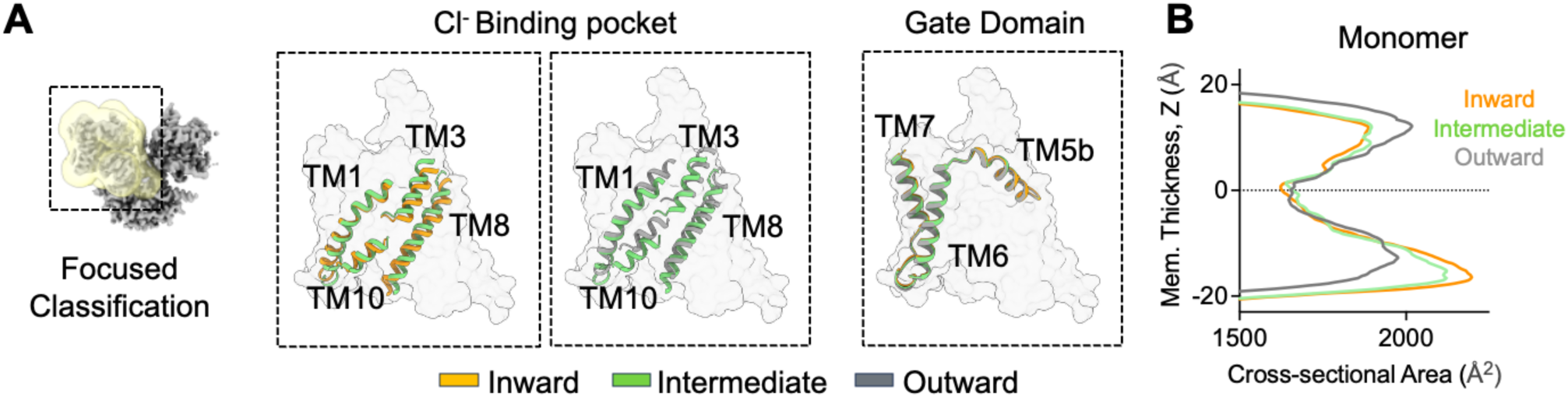
Focused 3D-classification of individual zebrafish SLC26A5 monomers. A) Focused 3D-classification of individual transmembrane domains in brain lipid nanodiscs revealed inward-facing, intermediate, and outward facing conformations. The additional intermediate conformation is mainly characterized by subtle movement of TMs 1,3 & 8 as compared to the inward-facing conformation. B) Comparison of the intra-membrane cross-sectional area of individual monomers in the three different conformations. Monomers were aligned based on residues 470-505.

**Fig. S13:**
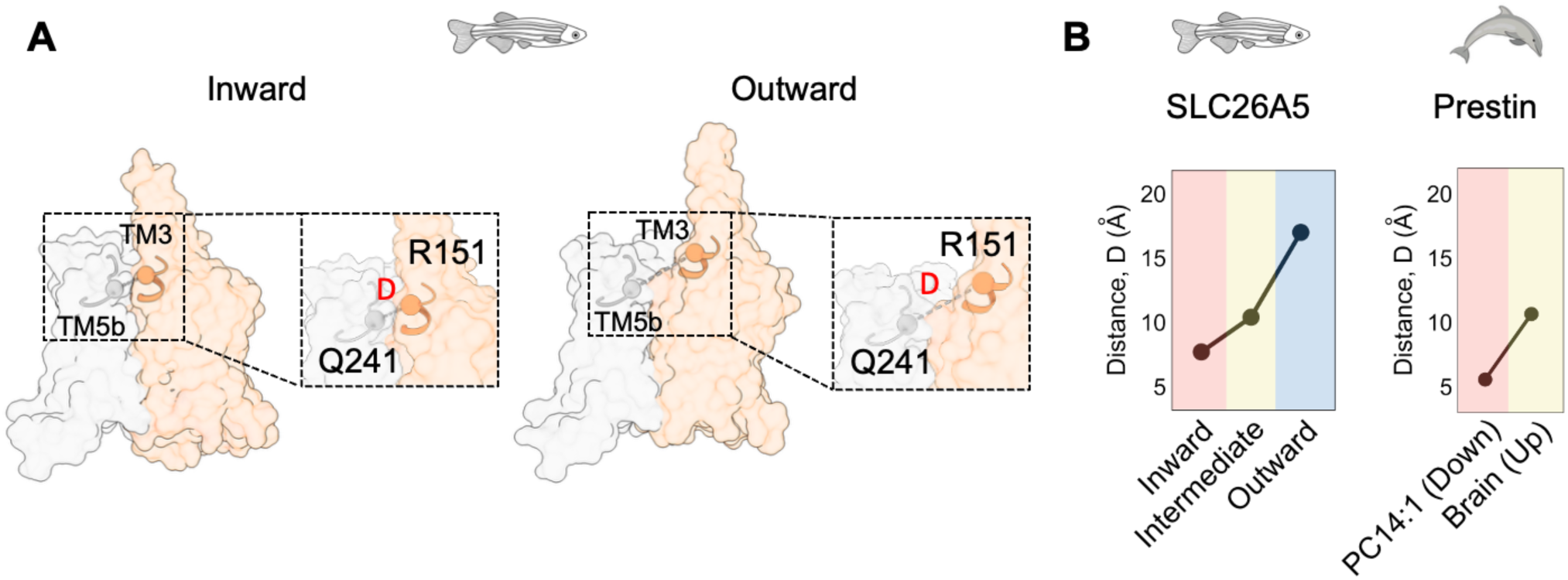
Extracellular accessibility of the anion binding site of zebrafish SLC26A5. A) Comparison of the distance (D) between the top of TM3 (α-carbon of R151) and TM5b (α-carbon of Q241) in inward vs outward facing states of zebrafish SLC26A5. In the outward facing conformation, TM3 moves away from TM5b, leading to the extracellular exposure of the anion binding site. B) Comparison of the distance between the α-carbon of R151 and α-carbon of Q241 in the different conformations of zebrafish SLC26A5 (left) and dolphin prestin (right). The corresponding residues for dolphin prestin are R150 and S238.

## Notes

### Competing Interest Statement

The authors have declared no competing interest.

### Summary of Updates

Updated refeneces, and reorganizing figures 1-4

